# A bifunctional kinase-phosphatase module integrates mitotic checkpoint and error-correction signalling to ensure mitotic fidelity

**DOI:** 10.1101/2022.05.22.492960

**Authors:** Andrea Corno, Marilia H Cordeiro, Lindsey A Allan, Qian Wei, Elena Harrington, Richard J Smith, Adrian T. Saurin

**Affiliations:** Division of Cellular and Systems Medicine, School of Medicine, University of Dundee, UK. DD1 9SY

## Abstract

Two major mechanisms have evolved to safeguard genome stability during mitosis: the mitotic checkpoint delays mitosis until all chromosomes have attached to microtubules, and the kinetochore-microtubule error-correction pathway keeps this attachment process free from errors. We demonstrate here that the optimal strength and dynamics of both processes is set by a kinase-phosphatase pair (PLK1-PP2A) that engage in negative feedback on the BUB complex. Uncoupling this homeostatic feedback to skew the balance towards PLK1 produces a strong checkpoint, weak microtubule attachments, and mitotic delays. Conversely, skewing the balance towards PP2A causes a weak checkpoint, strong microtubule attachments, and chromosome segregation errors. The number of MELT motifs on the KNL1 signalling scaffold sets the optimal levels of each enzyme, because engineering KNL1 to recruit too many BUB complexes increases KNL1-PLK1/PP2A levels, and enhances checkpoint/microtubule attachment strength. In contrast, recruiting too few BUB complexes lowers KNL1-PLK1/PP2A, and decreases checkpoint/microtubule attachment strength. Both of these situations are associated with chromosome segregation errors. Together, these data demonstrate how a single bifunctional kinase-phosphatase module integrates two major mitotic processes to help preserve genome stability.

## INTRODUCTION

Two key mitotic surveillance pathways have evolved to preserve genome stability by safeguarding the chromosome segregation process. The mitotic checkpoint (also known as the spindle assembly checkpoint or SAC) prevents mitotic exit until all chromosomes have attached to microtubules via the kinetochore (Lara-Gonzalez *et al*, 2021). The error-correction pathway continually monitors and corrects these attachments to ensure they remain free from errors (Lampson & Grishchuk, 2017). Both of these processes are regulated by dynamic phosphorylation events at the kinetochore (Saurin, 2018; Vallardi *et al*, 2017).

The SAC is activated by the phosphorylation of MELT repeats on the kinetochore signalling scaffold KNL1 (London *et al*, 2012; Shepperd *et al*, 2012; Yamagishi *et al*, 2012). These phosphorylated MELT repeats recruit the BUB complex to kinetochores, which act as platform for the assembly of an inhibitory complex that can block mitotic exit: termed the mitotic checkpoint complex or MCC (Lara-Gonzalez *et al*., 2021). Phosphorylation is also needed on multiple MCC components to drive complex assembly (Faesen *et al*, 2017; Ji *et al*, 2018; Ji *et al*, 2017; Qian *et al*, 2017; Zhang *et al*, 2017), and when MCC is generated, it is released from kinetochores to inhibit the anaphase promoting complex/cyclosome (APC/C); a large E3 ubiquitin ligase that otherwise induces chromosome segregation and mitotic exit by degrading securin and cyclin B (Lara-Gonzalez *et al*., 2021). The key phosphorylation events that drive MCC formation at kinetochores must be dynamic (i.e. responsive to change), because as soon as microtubule attach, MCC assembly must be rapidly shut down on KNL1. The kinetochore phosphatases PP2A-B56 and PP1, which bind to the BUB complex and KNL1 respectively, are crucial for dephosphorylating key sites to allow this rapid SAC silencing (Cordeiro *et al*, 2020; Espert *et al*, 2014; Espeut *et al*, 2012; London *et al*., 2012; Meadows *et al*, 2011; Nijenhuis *et al*, 2014; Rosenberg *et al*, 2011). This ultimately helps to ensure mitotic exit can occur within seconds after the last kinetochore attaches to microtubules.

The error-correction process also critically relies on rapidly switching phospho-sites, but in this case, the phosphorylation events are inhibitory to the microtubule attachment process (Saurin, 2018). That is because they are located on the microtubule attachment interface, including on multiple residues in the N-terminal tail of NDC80, where they electrostatically interfere with microtubule binding (Wimbish & DeLuca, 2020). The purpose of these phosphorylations is to help to correct attachment errors, and they achieve this by responding differently depending on the type of microtubule attachments that form. If the attachments are the correct amphitelic configuration - i.e. each sister kinetochore is attached to opposite spindle poles - then this exerts pulling force across the kinetochores. This “tension” is sensed, by a still poorly understood pathway, and the inhibitory phosphorylation sites on the attachment interface are dephosphorylated to stabilise microtubule binding. If, however, tension is not generated because the attachments are incorrect, then phospho-signals persist and those faulty attachments are destabilised, thus freeing the kinetochore to try again to form the correct amphitelic configuration (Lampson & Grishchuk, 2017; McVey *et al*, 2021). Rapidly responsive phospho-switching is crucial here too, because if phosphatases cannot dephosphorylate these sites quickly following tension, then even the correct microtubule attachments are destabilised and mitotic progression is delayed or prevented. That is essentially the phenotype observed when PP2A-B56 is removed from the BUB complex (Kruse *et al*, 2013; Suijkerbuijk *et al*, 2012; Xu *et al*, 2013), demonstrating that this phosphatase complex plays a crucial role in error-correction, as well as in SAC silencing. Why the same phosphatase complex is used for both process is still not clearly understood.

Although these two key mitotic surveillance pathways differ in their mechanism of action, they are united by a common requirement for their phosphorylation events to be both robust and dynamic. They must be robust, because if phosphorylation is too low, then the checkpoint is compromised and microtubule attachment errors are allowed to persist. The net result is a loss of mitotic fidelity as chromosome segregation errors occur, such as the type commonly seen in tumour cells with chromosomal instability (CIN) (Thompson *et al*, 2010). They must also be dynamic or responsive, because if phosphorylation events cannot change states quickly when microtubules bind and kinase activity is switched off, then attachments cannot be stabilised and the SAC cannot be extinguished. This prevents mitotic progression and eventually results in loss of chromosome segregation fidelity; for example, due to mitotic slippage or premature loss of sister chromatid cohesion. How kinetochores manage to balance these competing constraints of robustness and responsiveness remains unclear. What is clear, however, is that they manage to do this very efficiently because both processes are initially strong at kinetochores, but yet they are still able to rapidly shut down within seconds following a microtubule attachment (Clute & Pines, 1999; Dick & Gerlich, 2013). Understanding how they manage to achieve this could be crucial in the context of cancer, when robustness is compromised to produce the type of chromosome segregation errors that fuel tumour evolution.

We show here that human cells achieve the optimal trade-off between robustness and responsiveness, by using a kinase-phosphatase pair (PLK1-PP2A) that engage in homeostatic feedback on the BUB complex. This feedback loop balances the levels of a SAC activating kinase (PLK1) and a microtubule stabilising phosphatase (PP2A), such that both processes can function optimally to ensure mitotic fidelity. Together, this work demonstrates how a single bifunctional kinase-phosphatase module has evolved to safeguard chromosome segregation by integrating two key mitotic processes.

## RESULTS

### PLK1 and PP2A engage in an intramolecular negative feedback loop on the BUB complex

PP2A-B56 regulates both SAC silencing and KT-MT attachments by associating with the MCC component BUBR1 complex (Kruse *et al*., 2013; Suijkerbuijk *et al*., 2012; Xu *et al*., 2013), which localises to phosphorylated MELT repeats on KNL1 by binding to BUB1 and BUB3 (this BUB1-BUB3/BUB3-BUBR1 heterodimer is hereafter referred to as the BUB complex) (London *et al*., 2012; Overlack *et al*, 2015; Primorac *et al*, 2013; Shepperd *et al*., 2012; Vleugel *et al*, 2013; Yamagishi *et al*., 2012; Zhang *et al*, 2014). We demonstrated recently that PP2A-B56 controls SAC silencing principally by removing PLK1 from its phospho-binding motifs on BUBR1 (pT620) and BUB1 (pT609) (Cordeiro *et al*., 2020). From these residues, PLK1 can promote SAC signalling by phosphorylating the MELT repeats in an auto-catalytic loop that induces more BUB complex recruitment. In addition to phosphorylating the MELT motifs, PLK1 also phosphorylates the KARD domain on Ser676 and 680 to enhance PP2A recruitment to kinetochores (Elowe *et al*, 2007; Kruse *et al*., 2013; Suijkerbuijk *et al*., 2012; Wang *et al*, 2016a; Wang *et al*, 2016b). To examine if PLK1 phosphorylates the KARD from its binding site on BUBR1, we analysed KARD phosphorylation in BUBR1^WT^ and BUBR1^T620A^ (hereafter referred to as BUBR1^ΔPLK1^) cells. Note, all mutant experiments were performed after knockdown and replacement of the endogenous gene, unless stated otherwise.

Figures 1A and B show that S676 and T680 phosphorylation is reduced in BUBR1^ΔPLK1^ cells, demonstrating that local PLK1 recruitment is important for KARD phosphorylation. In contrast, S670 - a CDK1 site that also enhances PP2A-B56 binding (Kruse *et al*., 2013; Suijkerbuijk *et al*., 2012; Wang *et al*., 2016a; Wang *et al*., 2016b) - is unaffected by local PLK1 recruitment (Figure 1C). The PLK1 sites in the KARD are crucial for PP2A-B56 binding because BUBR1^ΔPLK1^ reduces B56γ at kinetochores to a similar extent as observed following deletion of the KARD domain (hereafter referred to a BUBR1^ΔPP2A(ΔK)^ - Figure 1D). Catalytic activity of PLK1 is crucial for these effects because a 30-min incubation with the PLK1 inhibitor, BI-2536 (Lenart *et al*, 2007), reduces pS676, pS680 and PP2A-B56 levels, similarly to BUBR1^ΔPLK1^ mutation (Figure S1). A crucial role for PLK1-mediated phosphorylation of the KARD is reinforced by the fact that the PLK1-regulated S676 is completely conserved in the KARD of BUBR1, and in the ancestral MADBUB homologue, throughout metazoa (as either a Ser or Thr residue; Figure 1I) (Cordeiro *et al*., 2020). Furthermore, the phospho-regulated PLK1 binding motif is almost always immediately adjacent to the KARD in BUBR1, or MADBUB homologues, and is often positioned around 50 amino acids prior to the KARD. Considering that the distance between these two binding domains was tightly conserved, we hypothesized that cross-regulation between PLK1 and PP2A occurs intra-molecularly.

**Figure 1.**
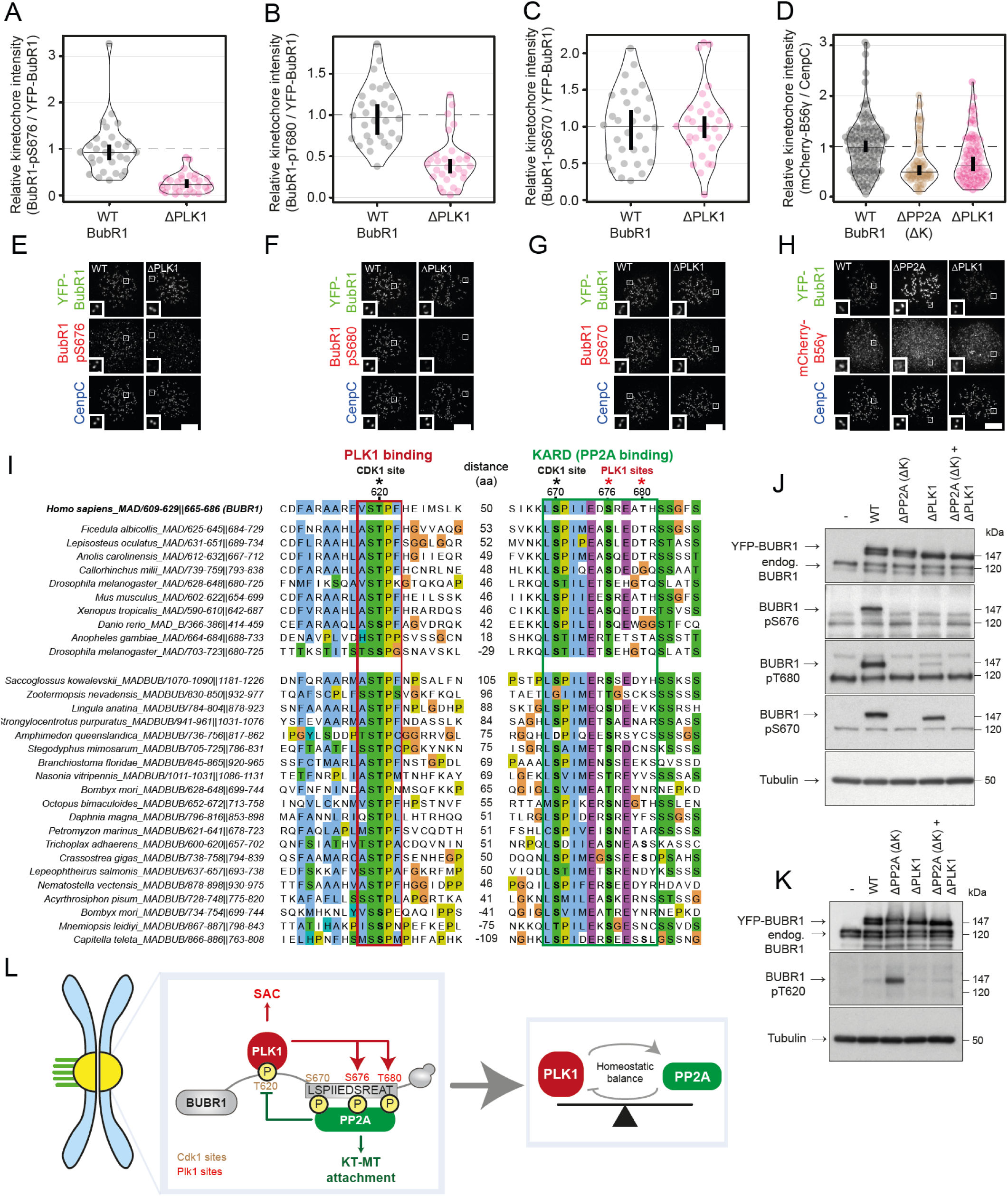
PLK1 and PP2A are engaged in an intramolecular negative feedback loop on the BUB complex. (A-D) Effects of preventing PLK1 binding to BUBR1 on levels of BUBR1-pS676 (A), BUBR1-pT680 (B), BUBR1-pS670 (C) and mCherry-B56γ (D) at unattached kinetochores, in nocodazole-arrested HeLa FRT cells expressing indicated YFP-tagged BUBR1 constructs (see Material and Methods for details). Kinetochore intensities from 30-80 cells, 3 to 5 experiments. **(E-H)** Example immunofluorescence images of the kinetochore quantifications shown in 1A-D. The insets show magnifications of the outlined regions. Scale bars: 5 μm. Inset size: 1.5 μm. **(I)** Alignment of PLK1 and PP2A binding region on MADBUB homologues in metazoa. The number of residues between PLK1 and PP2A binding is reported between the two alignments. **(J-K)** Mitotic HeLa FRT cells expressing indicated exogenous BUBR1 constructs were harvested and lysed. Lysates were then blotted with indicated antibodies. **(L)** Schematic illustrating how PLK1 and PP2A regulate each other binding to BUBR1. Kinetochore intensities are normalised to BUBR1 WT control (A-D). Violin plots show the distributions of kinetochore intensities between cells. For each violin plot, each dot represents an individual cell, the horizontal line represents the median, while the vertical one the 95% CI of the median, which can be used for statistical comparison of different conditions (see Materials and Methods).

To test this, we examined cross talk between endogenous BUBR1 and BUBR1 mutants that were unable to bind to either PLK1 (BUBR1^ΔPLK1^) or PP2A (BUBR1^ΔPP2A(ΔK)^) (i.e. by expressing mutants without knocking down the endogenous BUBR1). Figure 1J demonstrates that phosphorylation of S676 and T680 is reduced on YFP-tagged BUBR1^ΔPLK1^, but these sites remain largely unaltered on the endogenous BUBR1 protein, which is present in the same cells at similar levels. Similarly, T620 phosphorylation is only increased on YFP-BUBR1^ΔPP2A(ΔK)^, and not on the endogenous BUBR1^WT^ protein that is also present in the same cells (Figure 1K). Therefore, PLK1 and PP2A-B56 are engaged in an intra-molecular negative feedback loop on BUBR1, with PLK1 enhancing PP2A and PP2A decreasing PLK1. This homeostatic feedback loop likely acts to balance the kinetochore levels of the SAC kinase, PLK1, and microtubule stabilising phosphatase, PP2A-B56, such that their levels cannot increase too high or decrease too low (Figure 1L).

### The homeostatic balance between PLK1 and PP2A ensure strong and dynamic SAC and KT-MT attachment signalling

We hypothesised that balanced kinase-phosphatase recruitment to this region was crucial to allow optimal SAC and KT-MT attachment signalling. Therefore, to test this, we used mutants that we predicted would skew the balance towards either the kinase or the phosphatase (Figure 2A) (Smith *et al*, 2019). To create the phosphatase-dominant situation, B56γ was tethered to the C-terminus of BUBR1 in place of the KARD and pseudokinase domain (BUBR1^B56γ^). Note, that the only known function of the pseudokinase is to regulate KARD phosphorylation and therefore PP2A-B56 recruitment (Gama Braga *et al*, 2020). To create the analogous kinase-dominant situation, that KARD was deleted together with the pseudokinase domain by removing the entire C-terminus (BUBR1^ΔPP2A(ΔC)^). Figure 2B and C demonstrates that this system skews the balance as expected, because when the phosphatase is removed in BUBR1^ΔPP2A(ΔC)^ cells, BUBR1-pT620 and PLK1 levels increase at kinetochores, as shown previously for removal of just the KARD domain (BUBR1^ΔPP2A(ΔK)^) (Cordeiro *et al*., 2020). Conversely, when the phosphatase is fused in BUBR1^B56γ^ cells, BUBR1-pT620 and PLK1 levels are severely reduced at kinetochores. Note, this is equivalent to full removal of PLK1 from the BUB complex because kinetochores levels of PLK1 drop similarly to that seen with BUB1 depletion (Figures 2B, S2A-D).

**Figure 2.**
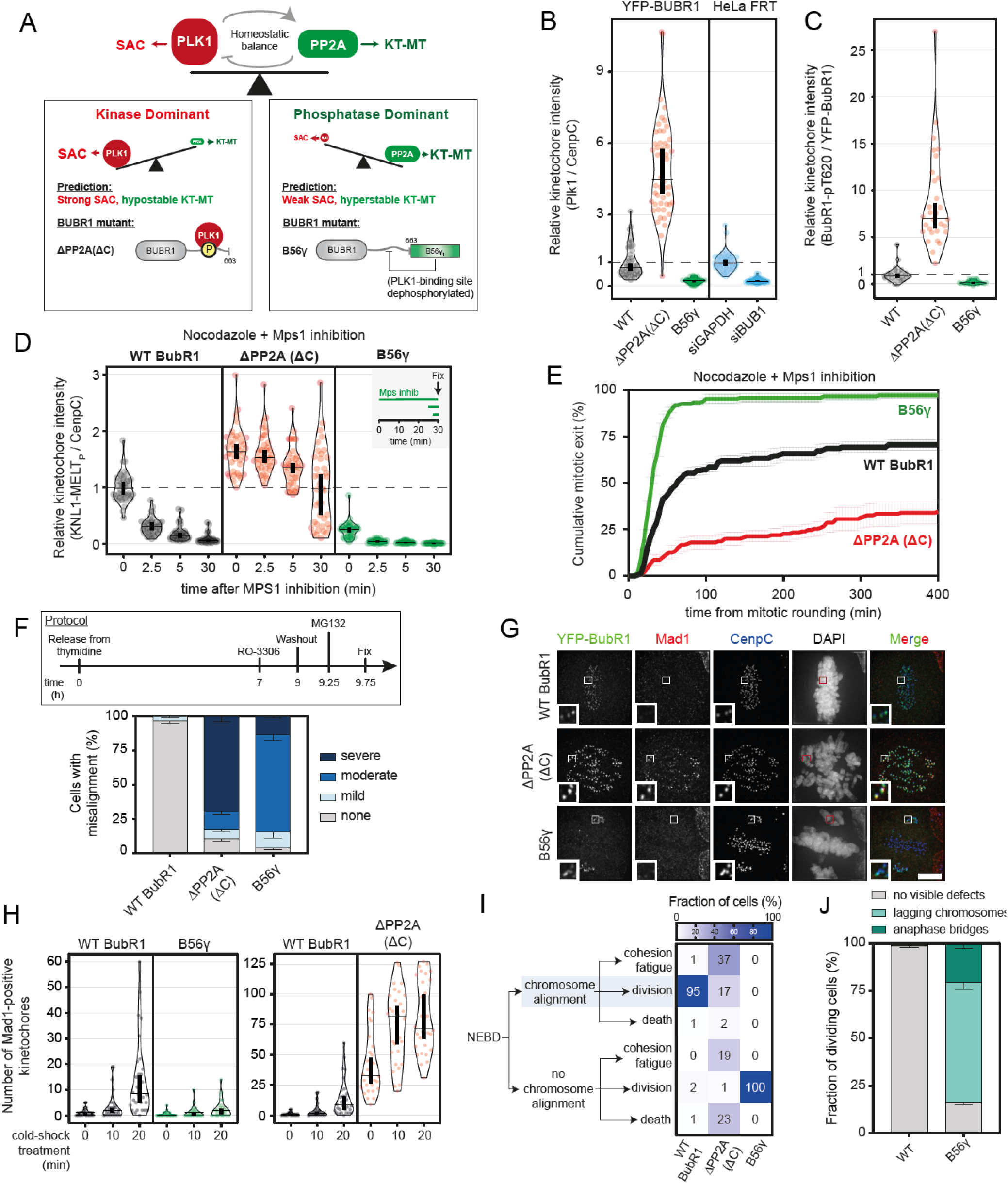
The homeostatic balance between PLK1 and PP2A ensures the strength and dynamics of the SAC and KT-MT attachments are optimal. **(A)** Schematic illustrating PLK1 and PP2A engaged in a homeostatic balance (on top) and the BUBR1 mutants used to lock the feedback loop on PLK1 (bottom-left panel) or on PP2A (bottom-right panel). **(B-C)** Effect of locking PLK1 or PP2A in the feedback loop on levels of PLK1 (B) and BUBR1-pT620 (C) at unattached kinetochores, in nocodazole-arrested HeLa FRT cells expressing the indicated BUBR1 mutants illustrated in (A) or treated with control or BUB1 siRNAs. Kinetochore intensities from 30-40 cells, 3 to 4 experiments. **(D-E)** Effects of locking PLK1 or PP2A in the feedback loop on KNL1-MELT dephosphorylation (D) and duration of mitotic arrest (E) in nocodazole-arrested cells treated with the MPS1 inhibitor AZ-3146 (2.5µM). Treatment with MG132 was included in (D) to prevent mitotic exit after the addition of the MPS1 inhibitor. (D) displays kinetochore intensities of 40 cells per condition, 4 experiments. (E) displays 50 cells per condition per experiment, 3 experiments. **(F-H)** Effects of locking PLK1 or PP2A in the feedback loop on chromosome alignment and kinetochore-microtubules attachments. (F) Top panel: protocol used to visualise chromosome misalignment in fixed samples (see Material and Methods for details). Bottom panel: graph showing mean frequencies (+/-SEM) of 3 experiments, 100 cells quantified per condition per experiment. (G) Example immunofluorescence images to show the presence of chromosome misalignments (DAPI) and the presence of unattached kinetochores (Mad1) in MG132-treated cells. The insets show magnifications of the outlined regions. Scale bars: 5 μm. Inset size: 1.5 µm. (H) The number of kinetochores positive for Mad1 was measured as a readout of unattached kinetochores. The measurement was performed on 30-40 **(Figure 2 continued)** cells from 4 experiments, during a cold-shock treatment to disrupt unstable kinetochore-microtubules attachments. Treatment with MG132 was included in (F,G and H) to prevent cells exiting mitosis. **(I-J)** Effects of locking PLK1 or PP2A in the feedback loop on the mitotic cell fate after Nuclear Envelope Breakdown (NEBD). The heatmap in (I) shows the mean frequencies of cell fates after NEBD in each condition – 3 experiments, 50 cells per condition per experiment (see also S2H). (J) The frequencies of errors in anaphase were scored for the dividing BUBR1 WT and B56γ cells showed in (I).

Given the role of PLK1 and PP2A in regulating SAC and KT-MT attachments, we hypothesised that the kinase-dominant situation would lead to a strong SAC and hypostable KT-MT attachments, whereas the phosphatase dominant situation would cause hyperstable KT-MT attachment and a weak SAC (Figure 2A). To test the effects of skewing the balance on SAC strength, we quantified KNL1-MELT phosphorylation and mitotic exit in cells treated with nocodazole and partial MPS1 inhibition, which are sensitized SAC assays that can report increases or decreases in SAC strength (Nijenhuis *et al*., 2014; Santaguida *et al*, 2010; Saurin *et al*, 2011). BUBR1^ΔPP2A(ΔC)^ enhanced KNL1-MELT phosphorylation and SAC strength, as expected (Figures 2D-E, S2E). This was shown previously in BUBR1^ΔPP2A(ΔK)^ cells (Nijenhuis *et al*., 2014), and the enhanced SAC strength was due to elevated PLK1 activity because PLK1 inhibition could completely rescue these effects (Cordeiro *et al*., 2020). Conversely, the opposite effects were seen in BUBR1^B56γ^ cells, which showed reductions in MELT phosphorylation and SAC strength (Figures 2D-E), consistent with the reduced PLK1 kinetochore levels in this situation (Figures 2B and S2F).

To examine if KT-MT attachments were similarly perturbed, we initially performed chromosome alignment assays when mitotic exit was blocked by MG132. This demonstrated severe misalignments in BUBR1^ΔPP2A(ΔC)^ cells (Figure 2F), as expected, and as shown previously in BUBR1^ΔPP2A(ΔK)^ cells (Kruse *et al*., 2013; Suijkerbuijk *et al*., 2012; Xu *et al*., 2013). Loss of PP2A-B56 in these situations is known to elevate phosphorylation sites on the KT-MT interface which destabilise end-on microtubule attachment, producing unattached kinetochores that stain positive for MAD1 (Smith *et al*., 2019). Conversely, when B56γ is fused to BUBR1 in BUBR1^B56γ^ cells, although there was also a strong misalignment phenotype (Figure 2F), in this case the unaligned kinetochores were MAD1-negative, indicating that they were stably attached to microtubules (Figure 2G). We hypothesised that these reflect hyperstable KT-MT attachment that were insensitive to the error-correction machinery. In agreement, kinetochores of BUBR1^B56γ^ cells were resistant to detachment following cold-shock treatment in comparison to BUBR1-WT cells (Figure 2H, left panel). This effect was inverted in BUBR1^ΔPP2A(ΔC)^ cells, which completely dissociated from microtubules within the 10 mins cold-shock treatment, as expected (Kruse *et al*., 2013; Suijkerbuijk *et al*., 2012) (Figure 2H).

In summary, a balanced recruitment of PLK1 and PP2A is needed to ensure optimal SAC and KT-MT regulation, and this balance is set by a bifunctional kinase-phosphatase module on BUBR1. Manipulating this module to shift the balance towards PLK1 results in a stronger SAC and unstable KT-MT attachments, whereas shifting the balance towards PP2A weakens the SAC and stabilises microtubules (Figure 2A). We hypothesised that these two abnormal states affect both the responsiveness and the robustness of the chromosome segregation process. That is because in the PLK1-dominant state, MELT phosphorylation is continually catalysed and phosphorylation sites that block KT-MT attachment are elevated (Smith *et al*., 2019), and most probably not turning over. Therefore, kinetochores are no longer responsive to MT attachment and the SAC signal persists. However, in the PP2A-dominant state, MELT phosphorylation is supressed and phosphorylation sites that block MT attachment are also low. Therefore, robustness of chromosome segregation is predicted to be lost as cells exit mitosis prematurely with incorrect attachments. Consistent with these predictions, live cell movies in BUBR1^B56γ^ cells demonstrated that these cells exit mitosis prematurely with many lagging chromosomes and anaphase bridges (Figures 2I-J, S2G-I, Movie S1 and S2). Note that this is not simply a weak checkpoint, because the prometaphase duration is actually extended in comparison to WT cells, indicating defective chromosome alignment as well (Figure S2J). In contrast, BUBR1^ΔPP2A(ΔC)^ cells arrest and struggle to align chromosomes, and even the cells that do visibly align chromosomes fail to switch-off the SAC and instead undergo cohesion fatigue (Figures 2I, S2G-J, Movie S1 and S3). In conclusion, a homeostatic feedback loop maintains an optimal balance of PLK1 and PP2A on the BUB complex, and this is crucial to ensure that SAC and KT-MT attachments signalling remains both strong and dynamic.

### The kinetochore levels of PLK1 and PP2A can be fined tuned by modulating the number of MELTs motifs on KNL1

The total level of PLK1 and PP2A at KNL1 are set by the number, sequence, and phosphorylation status of the MELT motifs. Human KNL1 contains up to 19 MELT motifs, although many of these have degenerated and lost key amino acids needed for BUB complex binding (Tromer *et al*, 2015). The BUB1 binding strength and the specific MELT sequences, as determined in (Vleugel *et al*, 2015) are shown in Figure 3A and B. We sought to modulate MELT numbers in a way that would allow BUB complex levels to be increased or decreased in a graded manner, thereby causing reciprocal changes to PLK1 and PP2A levels. MELT numbers have been reduced before in human KNL1, but this was achieved using artificial KNL1 fragments that also modified the total length and position of these motifs within KNL1 (Vleugel *et al*., 2013; Zhang *et al*., 2014). This could affect the ability of PLK1 and PP2A to signal from these artificial fragments, therefore we sought to change MELT number within the context of full length KNL1.

**Figure 3.**
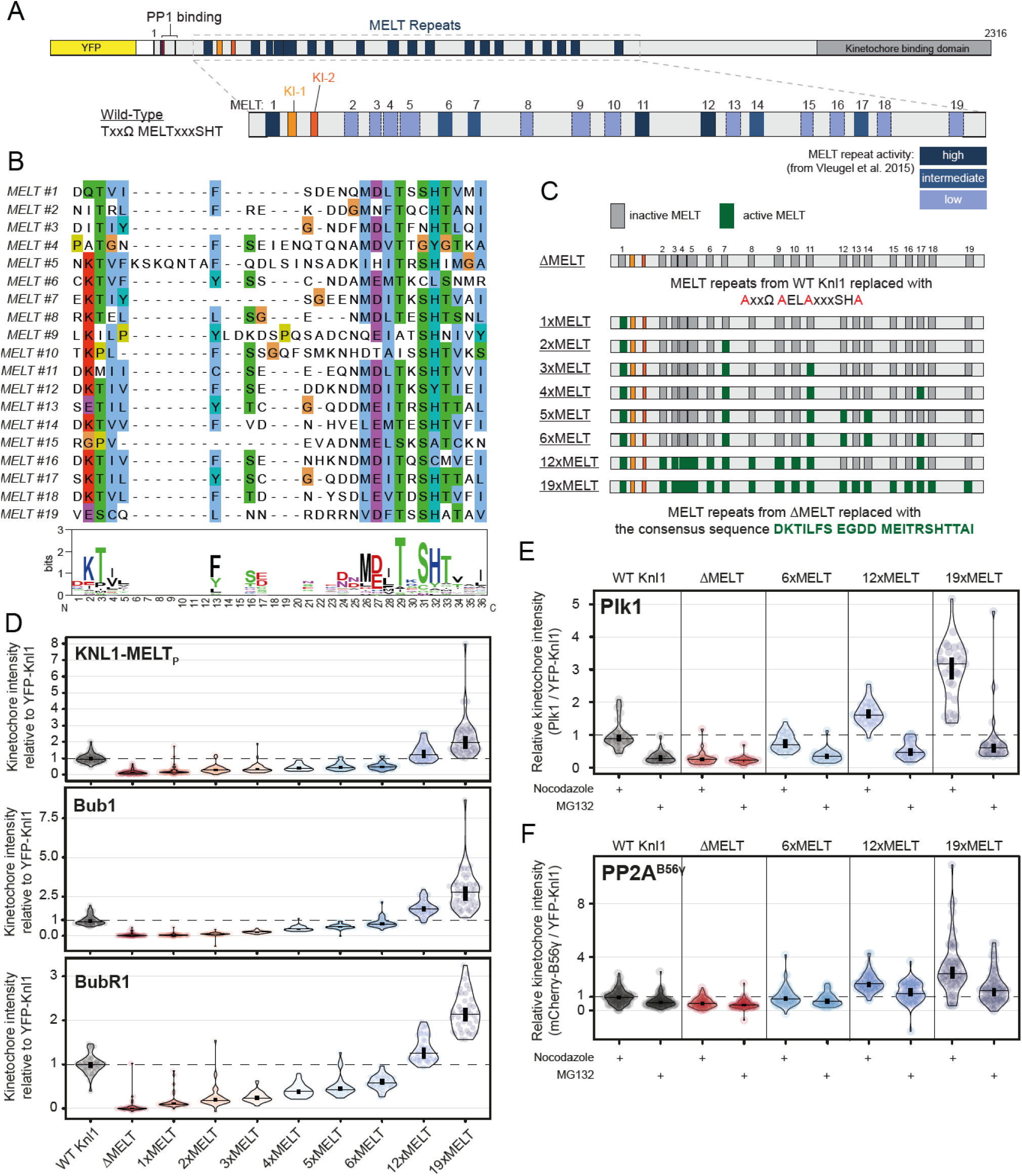
– KNL1 levels of the BUB complex, PLK1, and PP2A scale with the number of active MELT motifs. **(A)** Scheme of human KNL1 with N-terminal YFP-tag. MELT motifs are represented with different shades of blue, accordingly to the activity evaluated in (Vleugel *et al*., 2015). **(B)** Alignment of the 19 MELT motifs present in human Knl1. The weblogo represents the consensus sequence shared among all the motifs. **(C)** Schematic illustrating the KNL1 mutants created with a certain number of active MELTs (see Materials and Methods for details). **(D)** Levels of KNL1-pMELT, BUB1 and BUBR1 levels at unattached kinetochores, in nocodazole-arrested HeLa FRT cells expressing the KNL1 mutants shown in (C). Kinetochore intensities from 40-60 cells, 3 to 5 experiments. **(E, F)** Levels of PLK1 (E) and mCherry-B56γ (F) at kinetochores, in nocodazole or MG132-treated cells expressing the indicated KNL1-MELT mutants. Kinetochore intensities from 40-80 cells, 4 experiments. Kinetochore intensities are normalised to the WT KNL1 condition. Violin plots show the distributions of kinetochore intensities between cells. For each violin plot, each dot represents an individual cell, the horizontal line represents the median, while the vertical one the 95% CI of the median, which can be used for statistical comparison of different conditions (see Materials and Methods).

To do this, we first mutated key residues that are crucial for BUB complex binding on all possible MELT motifs within full length KNL1: referred to as KNL1^ΔMELT^ (Figure 3C) (Vleugel *et al*., 2015). Then we reintroduced an optimal MELT sequence, which our phospho-MELT antibody reacts with, into specific numbers of these MELT motifs ranging from 1 to 19 (Figure 3C). Figure 3D demonstrates that the kinetochore levels of phosphorylated MELT, BUB1, and BUBR1, are abolished in KNL1^ΔMELT^ cells, as expected. These levels are then increased in a graded manner as MELT number is increased, with 6xMELT recapitulating the closest to WT levels, and ≥12xMELTs producing artificially high BUB1/BUBR1 levels per KNL1 molecule. Note, total KNL1 kinetochore levels actually decreased in these situations, perhaps due to feedback between the BUB complex and KNL1 recruitment (Figure S3). We hypothesised that this system would also cause a similar graded reduction/increase in PLK1 and PP2A levels at KNL1, since these bind directly to BUBR1. This is shown in Figures 3E-F, with KNL1^ΔMELT^ reducing PLK1/PP2A, ≥ 12xMELTs increasing PLK1/PP2A, and 6xMELT recapitulating close to endogenous levels, regardless of KT-MT attachment status. In summary, this set of KNL1 mutants allow BUB complex levels to be precisely controlled at their native positions within full length KNL1. This can increase or decrease the levels of PLK1 *and* PP2A on each KNL1 molecule, crucially without modulating the feedback or skewing the balance between the two enzymes. We predicted that this would cause reciprocal changes in SAC and KT-MT attachment signalling such that too much kinase-phosphatase module would cause a strong SAC and hyperstable KT-MT attachment, and too little would cause a weak SAC and hypostable KT-MT attachments.

### The number of MELT motifs determines optimal PLK1 levels to ensure proper SAC regulation

We first assessed SAC signalling by examining mitotic arrest and KNL1-pMELT levels in nocodazole-arrested cells treated with 2.5 µM of the MPS1 inhibitor AZ-3146. We chose this dose because it causes a partial override of the SAC, therefore mutants that either strengthened or weakened the SAC response could be identified. Figure 4A demonstrates that the SAC is weakened in KNL1 mutants that contains ≤ 3xMELT motifs, strengthened with ≥ 5xMELT motifs, and indistinguishable from wild type in a mutant with 4xMELT motifs. We hypothesised that the strong SAC response was specifically due to elevated PLK1 at KNL1 (Figure 3E), which helps to amplify BUB levels by phosphorylating MELT motifs (Cordeiro *et al*., 2020). In agreement with this hypothesis, MELT phosphorylation and BUB1 levels were elevated in KNL1-19xMELT, and these remained high despite inhibition of MPS1 (Figure 4B,C). Furthermore, PLK1 activity was amplifying MELT signalling in this situation because PLK1 inhibition abolished BUB1 binding (Figure 4D) and allowed rapid mitotic exit (Figure 4E). Therefore, the number of MELT motifs is important to set the optimal kinetochore levels of PLK1 so that the SAC signal remains responsive to declining MPS1 levels.

**Figure 4.**
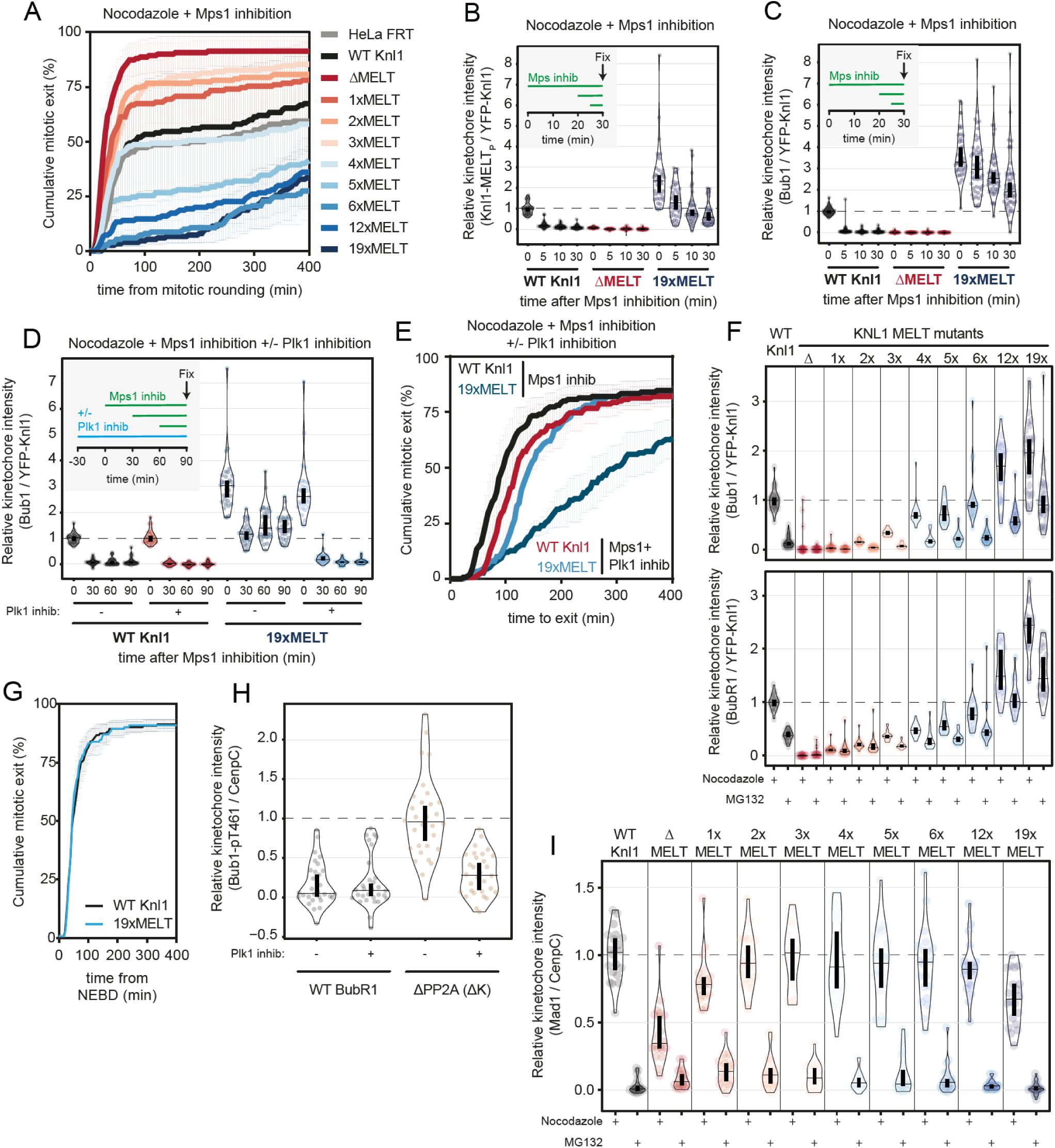
Modulating MELT number to fine-tune KNL1 levels of the PLK1/PP2A module impacts on SAC signalling. **(A-C)** Evaluation of the SAC signalling in KNL1-MELT mutants, in terms of the duration of the mitotic arrest (A), KNL1-MELT dephosphorylation (B) and BUB1 levels at unattached kinetochores (C), in nocodazole-arrested HeLa FRT cells expressing the indicated KNL1 mutants described in Figure 3C and treated with the MPS1 inhibitor AZ-3146 (2.5 µM). Treatment with MG132 was included in (B and C) to prevent mitotic exit after the addition of the MPS1 inhibitor. (A) shows the mean (+/-SEM) of 3 experiments, 50 cells per condition per experiment. (B, C) shows kinetochore intensities from 30 cells, 3 experiments. **(D, E)** Evaluating the contribution of PLK1 in sustaining the SAC signalling in WT KNL1 and 19xMELT cells, in terms of BUB1 levels at unattached kinetochores (D) and the duration of the mitotic arrest (E), in nocodazole-arrested cells expressing WT or 19xMELT KNL1 and treated with the MPS1 inhibitor AZ-3146 (2.5 µM) and with or without the PLK1 inhibitor BI-2536 (100 nM). (D) shows kinetochore intensities from 30 cells, 3 experiments. The graph in (E) shows mean (+/-SEM) of 3 experiments, 50 cells per condition per experiment. Treatment with MG132 was included in (D) to prevent mitotic exit after the addition of the MPS1 inhibitor. **(F)** Levels of BUB1 (top panel) and BUBR1 (bottom panel) at kinetochores, in nocodazole or MG132-treated cells expressing the KNL1 mutants described in Figure 3C. Kinetochore intensities from 10-30 cells, 2-3 experiments. **(G)** Cumulative distributions of the mitotic duration of WT and 19xMELT KNL1 cells during an unperturbed cell cycle. The graph reports the mean (+/-SEM) of 3 experiments, 42-50 cells per condition per experiment. **(H)** Levels of BUB1-pT461 at unattached kinetochores, in nocodazole-arrested HeLa FRT cells expressing BUBR1 WT or ΔPP2A (ΔK) – mutant in PP2A binding - and treated or not with the PLK1 inhibitor BI-2536 (100 nM). Kinetochore intensities from 30 cells, 3 experiments. **(I)** Levels of MAD1 at kinetochores, in nocodazole or MG132-treated cells expressing KNL1 mutants described in Figure 3C. Kinetochore intensities from 20 cells, 2 experiments. Kinetochore intensities are normalised to the WT KNL1 condition at time 0 in (B, C, D, F and I) or to the BUBR1^ΔPP2A(ΔK)^ untreated condition (H). Violin plots show the distributions of kinetochore intensities between cells. For each violin plot, each dot represents an individual cell, the horizontal line represents the median, while the vertical one the 95% CI of the median, which can be used for statistical comparison of different conditions (see Materials and Methods).

We next examined the effect of increasing MELT number on SAC silencing following KT-MT attachment/tension, by analysing BUB1/BUBR1 loss from KTs at metaphase. Figure 4F demonstrates that BUB1 and BUBR1 are better preserved at metaphase kinetochores when MELT number is increased, implying the elevate PLK1 in this situation (Figure 3E) is also able to better amplify MELT signalling on attached kinetochores. Mitotic duration is not extended under these conditions, however, demonstrating that the SAC is still silenced normally (Figure 4G). In addition to MELT phosphorylation, the SAC also relies on at least 2 other MPS1 phosphorylation sites on BUB1-pThr461) (Ji *et al*, 2017; Qian *et al*, 2017; Zhang *et al*, 2017) and MAD1-pThr716 (Faesen *et al*, 2017; Ji *et al*, 2018; Ji *et al*., 2017) to catalyse MCC assembly. We had shown previously that high PLK1 levels in BUBR1^ΔPP2A(ΔK)^ cells are unable to phosphorylate Mad1-Thr716 following Mps1 inhibition (Cordeiro *et al*. 2020), suggesting that, unlike the MELT motifs, this is a unique MPS1 site. Similar analysis on BUB1 demonstrates that PLK1 activity can contribute to Thr461 phosphorylation following PP2A removal (Figure 4H). The rise in pThr461 under these conditions has previously been attributed to reduced dephosphorylation (Qian *et al*., 2017), however, these data imply that enhanced phosphorylation by PLK1 is at least partially responsible. BUB1-pThr461 binds to MAD1 (Fischer *et al*, 2021; Zhang *et al*., 2017) therefore we analysed MAD1 levels at metaphase kinetochores. Figure 4I demonstrates that MAD1 is still effectively removed from kinetochore following KT-MT attachment, regardless of MELT numbers.

In summary, PLK1 is able to phosphorylate MELTs to recruit BUB1 and also phosphorylate BUB1 on Thr461 to recruit MAD1. This helps to amplify SAC signalling, especially when the number of MELT motifs are increased. We predict that increasing MELT numbers strengthens the positive feedback loop between PLK1 and BUB1 to prevent BUB dissociating from KNL1. However, even under these conditions, PLK1 activity alone cannot generate a SAC response, most probably because MAD1 is still removed from kinetochores at metaphase (Figure 4I). Interestingly, we noticed an abnormal recruitment of KNL1 to the midbody during anaphase in the 19xMELT mutant, along with BUB1, BUBR1 and PLK1 (Figure S4), suggesting that the KNL1 signalling platform cannot even disassemble at anaphase under these conditions. Therefore, the threshold at which PLK1 activity prevents the BUB complex from disassembling from KNL1 may have limited MELT expansion (Tromer *et al*., 2015).

### The number of MELT motifs determines optimal PP2A levels to ensure proper KT-MT attachment regulation

To examine how the KT-MT attachment process was affected by altering MELT levels, we performed live cell imaging to quantify chromosome segregation. Figures 5A and S5 demonstrates that in situations with less than 5 MELT motifs, chromosome alignment is perturbed and cells either die in mitosis, undergo cohesion fatigue, or divide with unaligned chromosomes. To analyse chromosome alignment more carefully under these conditions, we performed fixed assays in the presence of MG132. Figure 5B shows that KNL1^ΔMELT^ causes severe chromosome misalignments, as expected, given the PP2A-B56 binding to KNL1 is inhibited in this situation. These misalignments are progressively rescued by increasing MELT number until an optimal number of 6xMELT motifs, which appeared indistinguishable from wild type KNL1 (KNL1^WT^). Note, this is also the situation that rescued PP2A-B56 levels to close to wild type levels (Figure 3F). When MELT number is increased beyond 6xMELT, PP2A-B56 levels are elevated at KNL1 (Figure 3F), and there was a small but consistent increase in mild misalignments (Figure 5B). If this was due to hyperstable KT-MT attachments, as predicted, then these defects should become more apparent when cells are challenged to correct more KT-MT attachment errors. Therefore, we performed similar alignment assays after washout from an Eg5 inhibitor, STLC, to elevate KT-MT attachment errors (Kapoor *et al*, 2000). Chromosome alignment assays at different timepoints following washout demonstrated that the speed and total levels of chromosome alignment was impaired under conditions with 19xMELT motifs (Figure 5C). This is likely due to hyperstable KT-MT attachments because a short 10-min cold-shock treatment was less able to detach kinetochore fibres in KNL1-19xMELT cells, in comparison to KNL1-WT cells, as assayed by MAD1 accumulation to unattached kinetochores (Figure 5D). This was in contrast to KNL1^ΔMELT^ cells, which contained many unmatched kinetochores already prior to cold-shock, and these numbers increased, as expected, following the cold-shock treatment. Therefore, the number of MELT motifs is crucial for determining the stability of KT-MT attachments, most probably by setting the levels of PP2A-B56 on the KNL1/Mis12/NDC80 (KMN) network, which is a hub for KT-MT attachment regulation. This is important to prevent chromosome segregation errors in anaphase because the 19xMELT cells had elevated anaphase defects (Figure 5E).

**Figure 5.**
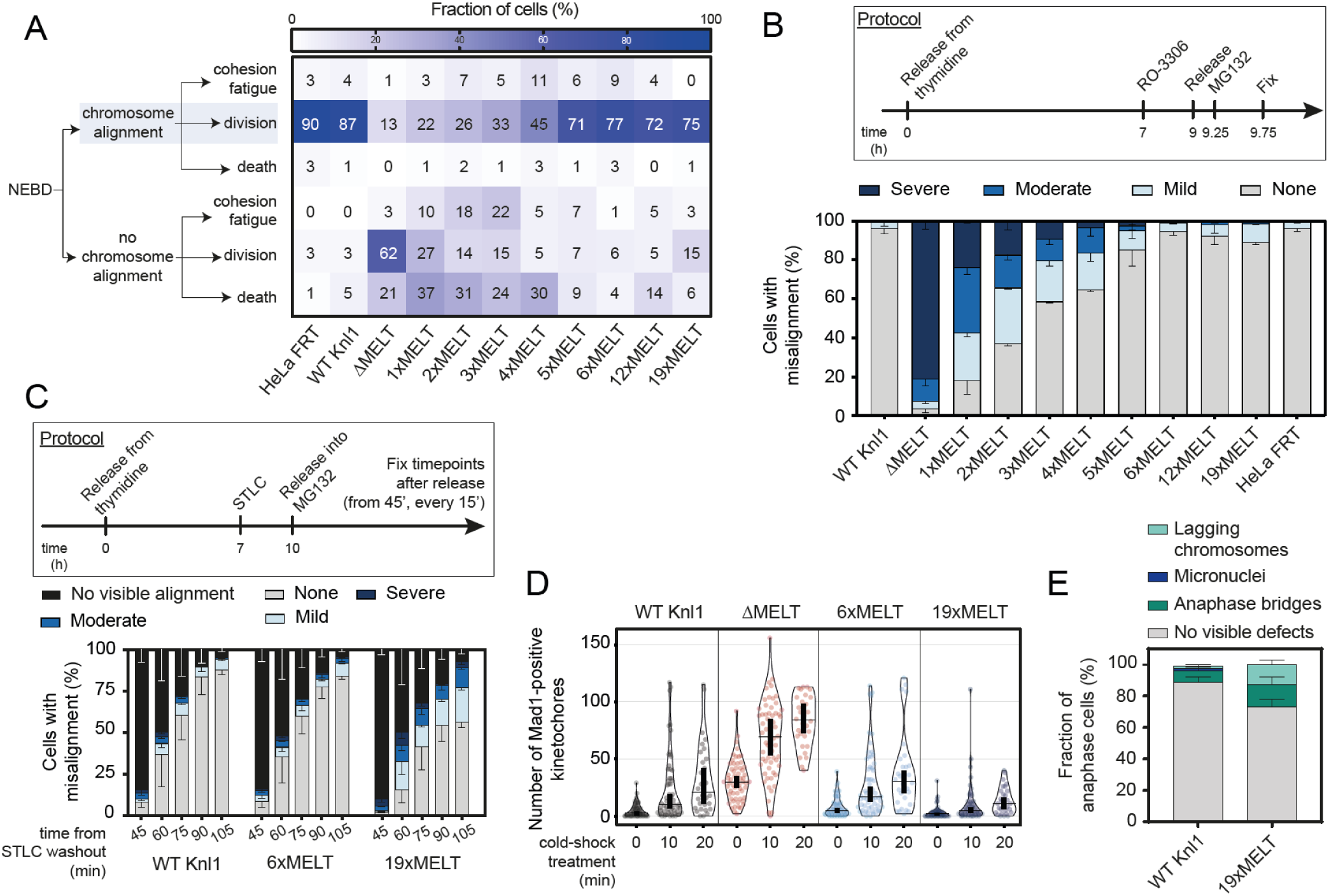
–Modulating MELT number to fine-tune KNL1 levels of the PLK1/PP2A module impacts on KT-MT attachments. **(A)** Mitotic cell fate after Nuclear Envelope Breakdown (NEBD) in cells expressing the KNL1-MELT mutants. The heatmap shows the mean frequencies of cell fates after NEBD in each condition – 3 experiments, 29-50 cells per condition per experiment (see also Figure S5). **(B-C)** Evaluating the effects on chromosome alignment in KNL1-MELT mutants. In both graphs, the top panel shows the protocol used to visualise chromosome misalignment in fixed samples (see Material and Methods for details), while the bottom panel reports mean frequencies (+/-SEM) of 3 experiment, 100 cells quantified per condition per experiment. **(D)** As a measure of unattached kinetochores, the number of the kinetochores positive for Mad1 recruitment was measured. The measurement was performed in cells expressing the indicated KNL1-MELT mutants, and during a cold-shock treatment to disrupt unstable kinetochore-microtubules attachments. 30-60 cells, 3 to 6 experiments. Treatment with MG132 was included in (B-D) to prevent mitotic exit. (E) Evaluation of the defects in chromosome segregation in WT and 19xMELT KNL1 cells. After a release from thymidine and incubation with STLC - as shown in (C) – cells were released into MG132-free media and fixed after 90’, to get an enrichment of anaphase cells (see Material and Methods for details). The graph shows mean (+/-SEM) of 3 experiments, 100 cells per condition per experiment. Violin plots show the number of kinetochores recruiting Mad1 between cells. For each violin plot, each dot represents an individual cell, the horizontal line represents the median, while the vertical one the 95% CI of the median, which can be used for statistical comparison of different conditions (see Materials and Methods).

## DISCUSSION

Here we have characterised a bifunctional kinase-phosphatase module on the BUB complex that is crucial to integrate SAC and KT-MT attachment signalling. A homeostatic negative feedback loop between the kinase PLK1 and phosphatase PP2A-B56 is crucial to limit the levels of these enzymes at kinetochores. This ensures that PLK1 can amplify SAC signalling by phosphorylating MELT repeats, without locking the SAC signal on. Thus, the SAC signal can remain strong, but crucially, still responsive to declining MPS1 activity. In turn, PP2A-B56 can stabilise initial KT-MT attachments without hyperstabilising them and impeding the KT-MT error-correction process. Therefore, KT-MT attachments can remain dynamic and responsive to tension. The final model is described in Figure 6.

**Figure 6:**
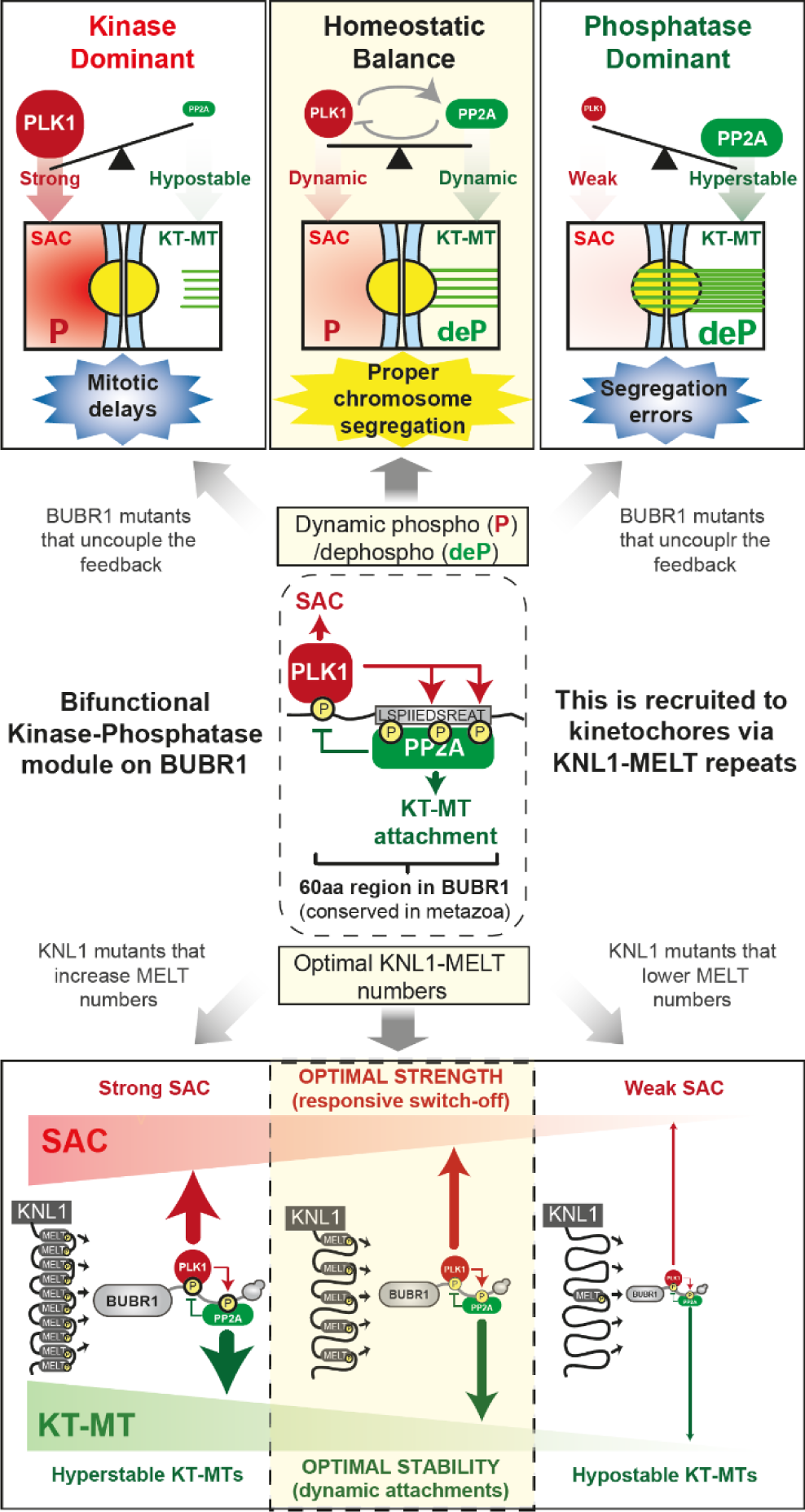
Schematic to illustrate how kinase-phosphatase coupling on the BUB complex integrates two mitotic processes to safeguard chromosome segregation. PLK1 and PP2A are engaged in a negative feedback loop on BUBR1 which ensure a homeostatic kinase-phosphatase balance and correct chromosome segregation. Top panel: Uncoupling the feedback alters the balance and causes chromosome segregation errors or mitotic delays. Bottom panel: the number of KNL1-MELT motifs sets the correct kinase-phosphatase levels to ensure optimal SAC strength and KT-MT stability.

The PLK1/PP2A module described here is recruited to kinetochore via the MELT motifs on KNL1, which is an important signalling hub for mitotic regulation (Ghongane *et al*, 2014). Previous studies on truncated versions of human KNL1 showed that the number of MELT repeats sets the kinetochore levels of the BUB complex and influences SAC signalling and chromosome alignment (Vleugel *et al*., 2013; Zhang *et al*., 2014). We expand on this here, by examining SAC signalling/strength, and chromosome alignment/KT-MT stability in a range of MELT mutants that span less than, and crucially, more than the optimal number. Importantly, this analysis was also performed within the context of full length KNL1. In agreement with previous studies, our data shows that > 3 MELT motifs are required for normal mitotic progression, but in contrast, we demonstrate that this relates to both SAC strength (Figure 4A) and KT-MT stability (Figure 5A). The ability of a KNL1-NC fusion containing just the first MELT motif to fully support the SAC in the Vleugel et al study, may be related to the artificially shortened KNL1, because within the context of full length KNL1, the first MELT motif alone exhibits a weaker SAC response (Figure 4A). By carefully analysing the effect of increasing and decreasing MELT number, we are able to link MELT numbers to PLK1/PP2A levels, and to the strength and responsiveness of the SAC and KT-MTs. Interestingly, a previous study examining MELT number on full length Spc105/KNL1 in *budding yeast*, also concluded that the MELT numbers were important to balance SAC strength and responsiveness, but for completely different reasons (Roy *et al*, 2020). It is notable that the PLK1-PP2A module is not conserved in *budding yeast* (Cordeiro *et al*., 2020), therefore they may have evolved different mechanisms to achieve the same end goal. Given the number and sequence of KNL1-MELTs is high variability in metazoa – due to iterative cycles of expansion and diversification (Tromer et *al*. 2015) – we propose that this has allowed the levels of this dynamic kinase-phosphatase module to rapidly evolve.

An important feature of combining kinase and phosphatase together in one module is that it ensures the levels of PLK1 and PP2A are coupled at kinetochores, such that their activities rise and fall together on KNL1. This helps to protect against the type of catastrophic chromosome segregation errors that would result if their activities became unbalanced. For example, if PP2A is allowed to predominate at kinetochores, the KMN network is dephosphorylated, the SAC is weakened and KT-MT attachments are stabilised, causing chromosome missegregations, such as the type observed in tumours with chromosomal instability (CIN).

These are essentially the phenotypes observed when the negative feedback in uncoupled in BUBR1^B56γ^ cells. In contrast, if PLK1 is allowed to predominate at kinetochores, increased phosphorylation of the KMN network causes SAC strengthening, decreased KT-MT stability, and mitotic delays. This occurs when feedback from PP2A-B56 is removed in BUBR1^ΔPP2A^ cells, resulting in decrease responsiveness and mitotic arrest. This can lead to either death in mitosis, mitotic slippage, or subsequent G1-arrest due to activation of the p53-dependent mitotic timer (Lambrus & Holland, 2017). In summary, the bifunctional kinase-phosphatase modules on the BUB complex integrates SAC and KT-MT attachment signalling to provide an optimal trade-off between a strong and responsive SAC, and stable and dynamic KT-MT attachments (Figure 6).

Considering this module is important to safeguard chromosome segregation, it is important to consider whether it could become altered or unbalanced to cause diseases, such as cancer. One interesting syndrome in this regard is mosaic-variegated aneuploidy (MVA): a rare childhood disorder characterised by chromosome missegregations and random aneuploidies in various somatic tissues, which leads to a diverse array of clinical features, including congenital abnormalities, developmental delays, and a predisposition to childhood cancers. Although 3 genes have been implicated in MVA syndrome, the majority of patients have mono or biallelic mutations in the BUB1B gene, which encodes for BUBR1 (Hanks *et al*, 2004; Yost *et al*, 2017). The primary effect of these mutations is to reduce BUBR1 protein level, which causes chromosome segregation defects due to the well-established roles of BUBR1 in the SAC and KT-MT attachment regulation (Suijkerbuijk *et al*, 2010). We propose that a major contributing factor to these segregation errors is reduced PLK1-PP2A recruitment to kinetochores, which as our work predicts, would weaken the SAC and KT-MT attachments.

Loss of BUBR1 protein has also been linked to age-related senescence in animal models and humans (Baker *et al*, 2004; Wijshake *et al*, 2012; Yang *et al*, 2017), and elevating BUBR1 expression can protect against many natural features of aging in mice (Baker *et al*, 2013). Similarly, we predict that lack of PLK1-PP2A recruitment to kinetochore could accelerate aging by causing chromosome missegregations and aneuploidies that drive cells into senescence. It will be interesting in future to determine whether the balance of PLK1/PP2A, rather than changes to steady-state levels, could be altered at kinetochores to drive cancer and/or ageing. There are many different protein that bind and inhibit PP2A-B56, some of which function at kinetochores (Asai *et al*, 2019; Asai *et al*, 2020; Porter *et al*, 2013), therefore release of these inhibitory pathways could perhaps tip the balance in favour of PP2A to weaken the SAC and stabilise KT-MTs in tumours cells with CIN.

Finally, kinases and phosphatases work together in many different ways to control the amplitude, localisation, timing and shape of phosphorylation signals (Gelens *et al*, 2018). In addition to that, they can also cooperate on individual molecules to drive cycles of phosphorylation and dephosphorylation to impart unique properties onto a signal response, such as robustness and responsiveness (Gelens & Saurin, 2018). For example, such “futile cycles” are used in EGFR signalling to provide a response that can rapidly change states (Kleiman *et al*, 2011). It is possible that these phosphorylation-dephosphorylation dynamics are crucial at kinetochores to allow signals to change state quickly upon microtubule attachment/tension, as discussed above. However, it is also possible that the picture becomes more complex if one considers what happens on each individual molecule. For example, individual BUB complexes may flip between the PLK1 or PP2A-bound state, with phosphorylation/dephosphorylation reactions driving cycles of PLK1/PP2A binding and release, as discussed in (Gelens & Saurin, 2018). It is also possible that these cycles work on specific subsets of BUB complexes at specific times. For example, BUB1/BUB3 complex may recruit PLK1 (via BUB1-pT609 (Cordeiro *et al*., 2020; Qi *et al*, 2006)) to amplify MELT phosphorylation, before BUBR1 association allows PP2A-B56 recruitment to remove PLK1, perhaps at a specific stage of MCC assembly. Understanding whether PLK1 and PP2A are recruited to specific BUB complexes at certain times, will be an important future goal. The answers to these questions may ultimately help to explain why this fascinating kinase-phosphatase module has been so well conserved throughout evolution.

## ACKNOWLEDGEMENTS

We thank staff at the Dundee Imaging Facility and the Genetic Core Services Unit. We also thank Stephen Taylor for providing the HeLa Flp-in cell line, Geert Kops and Mathieu Bollen for antibodies. This work was funded by a Cancer Research UK Programme Foundation Award to ATS (C47320/A21229 and C10988/A22566), which also funded AC, MHC, RS and LAA, and a Wellcome Investigator Award to ATS (222494/Z/21/Z), which funds AC and LAA. The authors declare no competing financial interests.

## AUTHOR CONTRIBUTIONS

ATS, MHC and AC conceived the study, designed the experiments, and interpreted the data. AC and MHC performed the majority of experiments, with help from LAA, QW, EH, and RJS. ATS supervised the study and wrote the manuscript together with AC, and with input from MHC.

## MATERIAL AND METHODS

### Cell culture

All cell lines used in this study were derived from HeLa Flp-in cells (a gift from S Taylor, University of Manchester, UK) (Tighe *et al*, 2008), which were authenticated by STR profiling (Eurofins). Cells were cultured in full growth media - DMEM supplemented with 9% FBS and 50 µg/ml penicillin/streptomycin. While doing fluorescence time-lapse analysis, cells were cultured in Leibovitz’s L-15 media (900 mg/L D+ Galactose, 5mM Sodium Pyruvate, no phenol red) supplemented with 9% FBS and 50µg/ml penicillin/streptomycin. Every 4-8 weeks cells were screened to ensure a mycoplasma free culture.

### Plasmids and cloning

pcDNA5-YFP-BUB1^WT^ and pcDNA5-YFP-BUBR1^WT^ expressing an N-terminally YFP-tagged and siRNA-resistant wild-type BUB1 or BUBR1 were described previously (Nijenhuis *et al*., 2014). pcDNA5-YFP-BUBR1^ΔPP2A(ΔK)^ (also called BUBR1^ΔKARD^), lacking amino acids 663-680 of BUBR1 was described previously (Nijenhuis *et al*., 2014). pcDNA5-YFP-BUBR1^ΔPP2A(ΔC)^, pcDNA5-YFP-BUBR1^B56γ^ (also called BUBR1^ΔCT^-B56γ_1_), pcDNA5-YFP-BUBR1^T620A^ (also called BUBR1^ΔPLK1^) and pcDNA5-YFP-BUBR1 ^ΔPP2A-T620A^ (also called BUBR1^ΔPP2A + ΔPLK1^) were described previously (Cordeiro *et al*., 2020; Smith *et al*., 2019).

All KNL1 constructs used in this study were derived from the plasmid pcDNA5-YFP-KNL1^WT^, which expresses an siRNA-resistant and N-terminally YFP-tagged wild-type KNL1 previously described (Smith *et al*., 2019). To create a full length KNL1 with variable number of active MELT motifs, we replaced the original motifs **T**xxΩ **M**EL**T**xxxSH**T** - which have variable lengths depending on their position in KNL1 (Vleugel *et al*., 2013) - with the amino acid sequence DK**T**ILFS EGDD **M**EI**T**RSH**T**TAI. This consensus sequence was designed based on conservation (Vleugel *et al*., 2013) as also represented in Figure 3B, strong affinity to recruit BUB complex (Vleugel *et al*., 2015) and optimal recognition using the pMELT antibody (pMELT-13/17, Thr 943 and Thr 1155, raised using the peptide MEIpTRSHTTALEC - (Nijenhuis *et al*., 2014). The nucleotide sequence of the active MELT repeats was varied as much as possible in the synthetised KNL1 constructs to avoid recombination of the plasmid during bacterial culture. To inactivate the MELT repeats and create KNL1^ΔMELT^, key threonines and methionines (or serines if present) in all 19 MELT motifs were mutated to alanine (**T**xxΩ **M**EL**T**xxxSH**T** was mutated to **A**xxΩ **A**EL**A**xxxSH**A**). All KNL1 constructs were subcloned by restriction cloning except when indicated. The pcDNA5-YFP-KNL1^ΔMELT^ and pcDNA5-YFP-KNL1^6xMELT^ (active MELT motifs at positions 1, 7, 11, 12, 14 and 17) were generated by inserting synthesized DNA fragments (Bio Basic Inc) in the backbone pcDNA5-YFP-KNL1^WT^ at restriction sites XmaI/Bsu36I and XhoI/BbvCI, respectively. To permit downstream cloning of different construct, silent mutations were introduced during DNA synthesis to insert extra restriction sites in the KNL1 constructs, including: XmaI before MELT-1, BssHII before MELT-6, HpaI before MELT-8, ApaI before MELT-12, XcmI before MELT-14 and Bsu36I before MELT-19. pcDNA5-YFP-KNL1^19xMELT^ was generated by Gibson assembly also using a synthesized DNA fragment (Bio Basic Inc) to insert extra MELT motifs in the pcDNA5-YFP-KNL1^6xMELT^ plasmid between PmlI and BbvCI sites. pcDNA5-YFP-KNL1^1xMELT-1^ (active MELT motif at positions 1), pcDNA5-YFP-KNL1^1xMELT-17^ (active MELT motif at positions 17), pcDNA5-YFP-KNL1^2xMELT^ (active MELT motifs at positions 1 and 7) and pcDNA5-YFP-KNL1^3xMELT^ (active MELT motifs at positions 1, 7 and 11) were created by inserting DNA fragments from pcDNA5-YFP-KNL1^6xMELT^ in the backbone pcDNA5-YFP-KNL1^ΔMELT^ using restriction sites XhoI/PmlI, AvrII/Bsu36I, XhoI/BlpI and XhoI/ApaI, respectively (KNL1^2xMELT^ generated using Gibson assembly with a fragment amplified by PCR). pcDNA5-YFP-KNL1^4xMELT^ (active MELT motifs at positions 1, 7, 11 and 17) and pcDNA5-YFP-KNL1^5xMELT^ (active MELT motifs at positions 1, 7, 11, 12 and 14) were created by inserting fragments from pcDNA5-YFP-KNL1^1xMELT-17^ or pcDNA5-YFP-KNL1 ^ΔMELT^ in the backbone pcDNA5-YFP-KNL1^6xMELT^ using ApaI/Bsu36I or AvrII/BbvCI sites, respectively. pcDNA5-YFP-KNL1^12xMELT^ (active MELT motifs at positions 1-11, and 17) was subcloned by replacing part of pcDNA5-YFP-KNL1^19xMELT^ (between ApaI/BbVCI restriction sites) with a fragment from pcDNA5-YFP-KNL1^1xMELT-17^. All plasmids were fully sequenced to verify the transgene was correct.

Cloning of pMESV_Ψ_-mCherry-B56γ_1_ plasmid has been described previously (Smith *et al*., 2019).

### Gene expression

HeLa Flp-in cells were stably generated to allows doxycycline-inducible expression of all constructs. Constructs were transfected with the relevant pcDNA5/FRT/TO plasmid and the Flp recombinase pOG44 (Thermo Fisher) using Fugene HD (Promega) according to the manufacturer’s instructions. Subsequently, cells were selected for stable integrants at the FRT locus using 200 µg/ml hygromycin B (Santa Cruz biotechnology) for at least 2 weeks. Cells expressing mCherry-B56γ_1_ were generated by viral-integration of pMESV_Ψ_-mCherry-B56γ_1_ construct into the genome of HeLa Flp-in cells, followed by puromycin selection (1mg/ml, Santa Cruz biotechnology). These cells were then used to stably express doxycycline-inducible KNL1^WT^, KNL1^ΔMELT^, KNL1^6xMELT^, KNL1^12xMELT^, KNL1^19xMELT^, BUBR1^WT^, BUBR1^ΔPLK1^ or BUBR1 ^ΔPP2A(ΔK)^, by following the same procedure described above.

### Gene knockdown

For all experiments involving re-expression in HeLa Flp-in cells, the endogenous mRNA was knocked down and replaced with an siRNA-resistant YFP-tagged mutant. The siRNAs used in this study were: siBUBR1 (5′-AGAUCCUGGCUAACUGUUC-3’), siBUB1 (5′-GAAUGUAAGCGUUCACGAA-3’) and siGAPDH (control siRNA: 5′-GUCAACGGAUUUGGUCGUA-3’). All siRNAs were synthesised with UU overhang (Sigma-Aldrich) and used at 20nM final concentration. Double stranded interference RNA was used to knockdown endogenous KNL1 (sense: 5’-GCAUGUAUCUCUUAAGGAAGAUGAA-3’; antisense: 5’-UUCAUCUUCCUUAAGAGAUACAUGCAU-3’) (Integrated DNA technologies) at a final concentration of 20 nM. All siRNAs/dsiRNAs were transfected using Lipofectamine® RNAiMAX Transfection Reagent (Thermo Fisher) according to the manufacturer’s instructions. After 16 h of knockdown, cells were arrested with thymidine (2mM, Sigma-Aldrich) for 24 h. Doxycycline (1µg/ml, Sigma-Aldrich) was used to induce expression of the BUBR1 and KNL1 constructs during and following the thymidine block. Cells were then released from thymidine block into full growth media supplemented with doxycycline and, when appropriate, nocodazole (3.3 µM, Sigma-Aldrich) for 5-7 hours for live imaging or 8.5 hours before processing for fixed analysis.

### Immunofluorescence

Cells plated on High Precision 1.5H 12-mm coverslips (Marienfeld) were fixed with 4% paraformaldehyde (PFA) in PBS for 10 min or pre-extracted with 0.1% Triton X-100 in PEM (100 mM PIPES, pH 6.8, 1 mM MgCl_2_ and 5 mM EGTA) for 1 minute before addition of 4% PFA for 10 minutes. After fixation, coverslips were washed with PBS and blocked with 3% BSA in PBS + 0.5% Triton X-100 for 30 min, incubated with primary antibodies overnight at 4°C, washed with PBS and incubated with secondary antibodies plus DAPI (4,6-diamidino2-phenylindole, Thermo Fisher) for an additional 2-4 hours at room temperature in the dark. Coverslips were washed with PBS and mounted on glass slides using ProLong antifade reagent (Molecular Probes). All images were acquired on a DeltaVision Core or Elite system equipped with a heated 37°C chamber, with a 100x/1.40 NA U Plan S Apochromat objective using softWoRx software (Applied precision). Images were acquired at 1×1 binning using a CoolSNAP HQ or HQ2 camera (Photometrics) and processed using softWorx software and ImageJ (National Institutes of Health). Mitotic cells were selected for imaging based on good expression of YFP at the kinetochore (KNL1) or cytoplasm (BUBR1 cells). All immunofluorescence images displayed are maximum intensity projections of deconvolved stacks and were chosen to closely represent the mean quantified data. Figure panels were creating using Omero (http://openmicroscopy.org).

The following primary antibodies (all diluted in 3% BSA in PBS) were used at the final concentration indicated: chicken anti-GFP (ab13970 from Abcam, 1:5000), rabbit anti-mCherry (GTX128508 from Genetex, 1:1000 – pre-extraction required to probe mCherry-B56γ_1_), guinea pig anti-CENP-C (PD030 from Caltag + Medsystems, 1:5000), rabbit anti-BUB1 (A300-373A from Bethyl, 1:1000), mouse anti-BUBR1 (05-898 from Millipore, 1:1000), rabbit anti-BUBR1 (A300-386A from Bethyl, 1:1000), rabbit anti-PLK1 (IHC-00071 from Bethyl, 1:1000), rabbit anti-BUBR1-pT680 (ab200061 from Abcam, 1:1000), mouse anti-α-tubulin (T5168-.2ML from Sigma-Aldrich, 1:5000) and mouse anti-MAD1 (MABE867 from Millipore, 1:1000).

The rabbit anti-pMELT-KNL1 antibody is directed against Thr 943 and Thr 1155 of human KNL1 (Nijenhuis *et al*., 2014) (1:1000 - gift from G. Kops, Hubrecht, NL). The rabbit anti-BUBR1-pT620 antibody was raised against phospho-Thr 620 of human BUBR1 using the peptide C-AARFVS[pT]PFHE (custom raised by Moravian, 1:1000, pre-extraction required) (Cordeiro *et al*., 2020). The rabbit anti-BUBR1-pS676 antibody was raised against phospho-S676 of human BUBR1 using the following peptide C-PIIED[pS]REATH (custom made by Biomatik, 1:200). The rabbit anti-BUBR1-pS670 antibody was raised against phosphor-Ser 670 of human BUBR1 (Nijenhuis *et al*., 2014) 1:1000. The rabbit anti-BUB1-pT461 is directed against Thr 461 of human BUB1 (1:500 – gift from M. Bollen, Leuven, BE) (Qian *et al*., 2017).

Secondary antibodies used were highly-cross absorbed goat anti-chicken Alexa Fluor 488 (A-11039), goat anti-rabbit Alexa Fluor 568 (A-11036), goat anti-mouse Alexa Fluor 488 (A-11029), goat anti-mouse Alexa Fluor 568 (A-11031), goat anti-guinea pig Alexa Fluor 647 (A-21450) or donkey anti-mouse Alexa Fluor 647 (A-31571) all used at 1:1000 (Thermo Fisher).

### Western Blotting

Protein lysates for immunoblot were prepared by harvesting mitotic cells, pelleting, and washing with cold PBS. After centrifuging samples at 1200 rpm for 3 min, pellets were lysed in ice cold RIPA buffer (50 mM Tris pH 8.0, 150 mM NaCL, 1% NP40, 0.5% sodium deoxycholate, 2mM EDTA pH 8.0, 0.1% SDS, 50 mM NaF and protease inhibitor cocktail) on ice for 20 min. Lysates were centrifuged at 13,000 g at 4°C for 10 min, followed by DC Protein Assay (Biorad) to estimate the concentration of each sample. Samples were then mixed with loading buffer to final concentrations of 62.5 mM Tris pH 6.8, 2.5% SDS, 10% glycerol, 5% β-mercaptoethanol and bromophenol blue. Samples were boiled, then run on 8% SDS-PAGE gels, and transferred onto PVDF. Blots were then blocked and incubated overnight at 4°C in primary antibody. Then membranes were washed in TBS with 0.1% Tween 20 (TBS-T), incubated in HRP-conjugated secondary antibody (BioRad) for 1 h at RT, washed in TBS-T and imaged with ECL.

The following primary antibodies (all diluted in 3% BSA in PBS) were used at the final concentration indicated: rabbit anti-BUBR1 (A300-386A from Bethyl, 1:1000 in 5% milk in TBS-T after blocking in 5% milk in TBS-T), rabbit anti-BUBR1-pT680 (ab200061 from Abcam, 1:1000 in 5% milk in TBS-T after blocking in 5% milk in TBS-T), rabbit anti-BUBR1-pT620 (custom raised by Moravian, 1:1000 in 5% BSA in TBS-T after blocking in 5% milk in TBS-T), rabbit anti-BUBR1-pS670 (Nijenhuis *et al*., 2014) (1:1000 in 5% milk in TBS-T after blocking in 5% milk in TBS-T)(Suijkerbuijk *et al*., 2012), and rabbit anti-BUBR1-pS676 (custom raised by Biomatik, and used at 1:750 in ReliaBLOT® Block – Bethyl labs - after blocking in ReliaBLOT® Block, as per manufacturer’s instructions), and mouse anti-α-tubulin (T5168-.2ML from Sigma-Aldrich, 1:5000).

### SAC strength assays

To measure SAC strength in live cells, cells were incubated in a 24-well plate in full growth media in a heated chamber (37°C and 5% CO_2_) and imaged with brightfield microscopy using a 10x/0.5 NA objective and a Hamamatsu ORCA-ER camera at 2×2 binning on a Zeiss Axiovert 200M, controlled by Micro-manager software (open source: https://micro-manager.org/) or a 20x/0.4 NA air objective and a CMOS Orca flash 4.0 camera at 4×4 binning on a Zeiss Axio Observer 7. Mitotic exit was defined by cells flattening down in the presence of nocodazole and MPS1 +/-PLK1 inhibitors. MPS1 was inhibited with AZ-3146 (2.5 µM, Sigma-Aldrich) shortly prior to imaging, with or without PLK1 inhibition with BI-2536 (100 nM, Selleckbio). In Figures 2E and 4A, cells entering in mitosis in the presence of AZ-3146 were analysed. In Figure 4E, cells arrested in mitosis at the time of AZ-3146 +/-BI-2536 treatment were analysed.

To measure SAC strength in fixed cells, nocodazole and MG132 (10 µM, Sigma-Aldrich) +/-BI-2536 100nM were added first for 30 minutes to ensure complete inhibition, followed by a time-course of AZ-3146 2.5μM +/-BI-2536 100nM in media containing nocodazole and MG132. Cells were fixed and analysed by immunofluorescence, probing for KNL1-pMELT or BUB1.

### Chromosome alignment assays

To observe live chromosome alignment and determine mitotic cell fates and timing in an unperturbed cell cycle, cells were plated in 8-well or 18-well chamber slides (ibidi), released from thymidine block for 5-6 hours, incubated with SiR-DNA far-red DNA probe (1:10000, Spirochrome; to prevent toxicity (Sen *et al*, 2018)) in L-15 media for 15min, rinsed and imaged every 4 minutes during 16 hours with a 40x/1.3 oil immersion objective or 40x/0.95 air objective using a Zeiss Axio Observer 7 with a CMOS Orca flash 4.0 camera at 4×4 binning, 10 z-stacks and a step size of 1.50 µm. Cells were selected for quantification based on good expression of YFP-tagged protein. Selected cells were scored based on the following mitotic events: cohesion fatigue, cell division or cell death following chromosome alignment or not. Dividing cells were also scored based on the type of chromosome segregation defect (no visible defects, anaphase bridges or lagging chromosomes).

To observe chromosome alignment in fixed-cell experiments – with the advantage of taking high resolution images and easily selecting many cells for further analysis - cells were released from thymidine block for 7 hours before being treated for 2 hours with RO-3306 (10μM, Tocris) – to synchronise cells at the G2/M boundary - or with a 3-hour treatment with STLC (10μM, Sigma-Aldrich) - to arrest cells in mitosis with monopolar spindles. Cells treated with RO-3306 were then washed three times and incubated for 15 minutes with full growth media before addition of MG132 to prevent mitotic exit, fixing cells 30’ after the addition of MG132. Cells treated with STLC were then washed three times and incubated in full growth media supplemented with MG132, fixing cells every 15’ from 45’ to 105’ after the addition of MG132. Fixed cells were stained as described above and imaged on a Zeiss Axio Observer with a CMOS Orca flash 4.0 camera at 4×4 binning, using a Plan-apochromat 20×/0.4 air objective. Cells with a good expression of YFP-tagged protein were scored based on the number of misaligned chromosomes as aligned (0 misaligned chromosomes), mild (1-2), moderate (3-5), severe (>6) or no visible alignment (for cells released from STLC in which a clear metaphase plate was not visible – either because of a monopolar spindle induced by the STLC treatment or because the metaphase plate was rotated). This protocol is important because mutants that cause a prolonged arrest can otherwise cause cohesion fatigue, which skews the alignment data.

To observe chromosome segregation defects in fixed-cell experiments, cells were treated with STLC as described above, but then released in MG132-free media to allow cells to proceed into anaphase. Cells were fixed 90’ after the STLC washout and imaged as describe above. Anaphase cells with a good expression of YFP-tagged protein were scored based on the type of chromosome segregation defect (no visible defects, anaphase bridges, lagging chromosomes or micronuclei).

To measure the proportion of unstable kinetochore-microtubules attachments, cells were treated with RO-3306 as described above in chromosome alignment experiments. After the RO-3306 washout followed by a 30-min treatment with MG132, the media was replaced with ice-cold L-15 supplemented with MG132 and cells were incubated in ice to induce cold-shock. Cells were then fixed 0, 10 and 20 mins after the temperature shift, and stained for Mad1 (i.e. marker of unattached kinetochores). The number of kinetochores unattached to the mitotic spindle was estimated by counting the number of the kinetochores positive for Mad1 recruitment.

### Comparison of kinetochore protein levels in prometaphase versus metaphase

To compare kinetochore protein levels in cells enriched in prometaphase versus metaphase, cells were treated with RO-3306 as described above in chromosome alignment experiments. After the RO-3306 washout, cells were treated with nocodazole or MG132 for 30’, to enrich cells in prometaphase and metaphase respectively. Cells were then fixed and stained as described above.

### Image quantification and statistical analysis

For quantification of kinetochore protein levels, images of similarly stained experiments were acquired with identical illumination settings and analysed using an ImageJ macro, as described previously (Saurin *et al*., 2011). Fluorescence intensities at kinetochores were normalised to CenpC (i.e. kinetochore marker). Measurements of phosphorylated BUBR1 were normalised to those of YFP-BUBR1, to avoid artificial fluctuation of signal resulting from variability in re-expression levels of YFP-BUBR1. The same principle was adopted in YFP-KNL1 cells to measure levels of BUB1, BUBR1, KNL1-pMELT, PLK1 and mCherry-B56γ.

Violin plots were produced using PlotsOfData - https://huygens.science.uva.nl/PlotsOfData/ (Postma & Goedhart, 2019). This allows the spread of data to be accurately visualised along with the 95% confidence intervals (thick vertical bars) calculated around the median (thin horizontal lines). This representation allows the statistical comparison between all treatments and timepoints because when the vertical bar of one condition does not overlap with one in another condition the difference between the medians is statistically significant (p<0.05).

### Analysis of PLK1 and PP2A binding motifs in BUBR1 throughout metazoa

The dataset from (Tromer *et al*, 2016) was used and annotated for the presence of PLK1 and PP2A binding sites in metazoa, as described previously (Cordeiro *et al*., 2020). Sequence alignments were generated using Jalview (Waterhouse *et al*, 2009) The consensus sequence for MELT motifs was created using WebLogo (Crooks *et al*, 2004); https://weblogo.berkeley.edu/logo.cgi).

## REFERENCES

Asai Y, Fukuchi K, Tanno Y, Koitabashi-Kiyozuka S, Kiyozuka T, Noda Y, Matsumura R, Koizumi T, Watanabe A, Nagata K et al (2019) Aurora B kinase activity is regulated by SET/TAF1 on Sgo2 at the inner centromere. J Cell Biol 218: 3223–3236

Asai Y, Matsumura R, Hasumi Y, Susumu H, Nagata K, Watanabe Y, Terada Y (2020) SET/TAF1 forms a distance-dependent feedback loop with Aurora B and Bub1 as a tension sensor at centromeres. Sci Rep 10: 15653

Baker DJ, Dawlaty MM, Wijshake T, Jeganathan KB, Malureanu L, van Ree JH, Crespo-Diaz R, Reyes S, Seaburg L, Shapiro V et al (2013) Increased expression of BubR1 protects against aneuploidy and cancer and extends healthy lifespan. Nat Cell Biol 15: 96–102

Baker DJ, Jeganathan KB, Cameron JD, Thompson M, Juneja S, Kopecka A, Kumar R, Jenkins RB, de Groen PC, Roche P et al (2004) BubR1 insufficiency causes early onset of aging-associated phenotypes and infertility in mice. Nat Genet 36: 744–749

Clute P, Pines J (1999) Temporal and spatial control of cyclin B1 destruction in metaphase. Nat Cell Biol 1: 82–87

Cordeiro MH, Smith RJ, Saurin AT (2020) Kinetochore phosphatases suppress autonomous Polo-like kinase 1 activity to control the mitotic checkpoint. J Cell Biol 219

Crooks GE, Hon G, Chandonia JM, Brenner SE (2004) WebLogo: a sequence logo generator. Genome research 14: 1188–1190

Dick AE, Gerlich DW (2013) Kinetic framework of spindle assembly checkpoint signalling. Nat Cell Biol 15: 1370–1377

Elowe S, Hummer S, Uldschmid A, Li X, Nigg EA (2007) Tension-sensitive Plk1 phosphorylation on BubR1 regulates the stability of kinetochore microtubule interactions. Genes Dev 21: 2205–2219

Espert A, Uluocak P, Bastos RN, Mangat D, Graab P, Gruneberg U (2014) PP2A-B56 opposes Mps1 phosphorylation of Knl1 and thereby promotes spindle assembly checkpoint silencing. J Cell Biol 206: 833–842

Espeut J, Cheerambathur DK, Krenning L, Oegema K, Desai A (2012) Microtubule binding by KNL-1 contributes to spindle checkpoint silencing at the kinetochore. J Cell Biol 196: 469–482

Gama Braga L, Cisneros AF, Mathieu MM, Clerc M, Garcia P, Lottin B, Garand C, Thebault P, Landry CR, Elowe S (2020) BUBR1 Pseudokinase Domain Promotes Kinetochore PP2A-B56 Recruitment, Spindle Checkpoint Silencing, and Chromosome Alignment. Cell reports 33: 108397

Gelens L, Qian J, Bollen M, Saurin AT (2018) The Importance of Kinase-Phosphatase Integration: Lessons from Mitosis. Trends Cell Biol 28: 6–21

Gelens L, Saurin AT (2018) Exploring the Function of Dynamic Phosphorylation-Dephosphorylation Cycles. Developmental cell 44: 659–663

Kapoor TM, Mayer TU, Coughlin ML, Mitchison TJ (2000) Probing spindle assembly mechanisms with monastrol, a small molecule inhibitor of the mitotic kinesin, Eg5. J Cell Biol 150: 975–988

Kleiman LB, Maiwald T, Conzelmann H, Lauffenburger DA, Sorger PK (2011) Rapid phospho-turnover by receptor tyrosine kinases impacts downstream signaling and drug binding. Molecular cell 43: 723–737

Kruse T, Zhang G, Larsen MS, Lischetti T, Streicher W, Kragh Nielsen T, Bjorn SP, Nilsson J (2013) Direct binding between BubR1 and B56-PP2A phosphatase complexes regulate mitotic progression. J Cell Sci 126: 1086–1092

Lampson MA, Grishchuk EL (2017) Mechanisms to Avoid and Correct Erroneous Kinetochore-Microtubule Attachments. Biology (Basel) 6

Lara-Gonzalez P, Pines J, Desai A (2021) Spindle assembly checkpoint activation and silencing at kinetochores. Semin Cell Dev Biol 117: 86–98

Lenart P, Petronczki M, Steegmaier M, Di Fiore B, Lipp JJ, Hoffmann M, Rettig WJ, Kraut N, Peters JM (2007) The small-molecule inhibitor BI 2536 reveals novel insights into mitotic roles of polo-like kinase 1. Curr Biol 17: 304–315

London N, Ceto S, Ranish JA, Biggins S (2012) Phosphoregulation of Spc105 by Mps1 and PP1 regulates Bub1 localization to kinetochores. Curr Biol 22: 900–906

McVey SL, Cosby JK, Nannas NJ (2021) Aurora B Tension Sensing Mechanisms in the Kinetochore Ensure Accurate Chromosome Segregation. Int J Mol Sci 22

Meadows JC, Shepperd LA, Vanoosthuyse V, Lancaster TC, Sochaj AM, Buttrick GJ, Hardwick KG, Millar JB (2011) Spindle checkpoint silencing requires association of PP1 to both Spc7 and kinesin-8 motors. Developmental cell 20: 739–750

Nijenhuis W, Vallardi G, Teixeira A, Kops GJ, Saurin AT (2014) Negative feedback at kinetochores underlies a responsive spindle checkpoint signal. Nat Cell Biol 16: 1257–1264

Overlack K, Primorac I, Vleugel M, Krenn V, Maffini S, Hoffmann I, Kops GJ, Musacchio A (2015) A molecular basis for the differential roles of Bub1 and BubR1 in the spindle assembly checkpoint. eLife 4: e05269

Porter IM, Schleicher K, Porter M, Swedlow JR (2013) Bod1 regulates protein phosphatase 2A at mitotic kinetochores. Nature communications 4: 2677

Postma M, Goedhart J (2019) PlotsOfData-A web app for visualizing data together with their summaries. PLoS Biol 17: e3000202

Primorac I, Weir JR, Chiroli E, Gross F, Hoffmann I, van Gerwen S, Ciliberto A, Musacchio A (2013) Bub3 reads phosphorylated MELT repeats to promote spindle assembly checkpoint signaling. eLife 2: e01030

Qi W, Tang Z, Yu H (2006) Phosphorylation- and polo-box-dependent binding of Plk1 to Bub1 is required for the kinetochore localization of Plk1. Mol Biol Cell 17: 3705–3716

Rosenberg JS, Cross FR, Funabiki H (2011) KNL1/Spc105 recruits PP1 to silence the spindle assembly checkpoint. Curr Biol 21: 942–947

Santaguida S, Tighe A, D’Alise AM, Taylor SS, Musacchio A (2010) Dissecting the role of MPS1 in chromosome biorientation and the spindle checkpoint through the small molecule inhibitor reversine. J Cell Biol 190: 73–87

Saurin AT (2018) Kinase and Phosphatase Cross-Talk at the Kinetochore. Front Cell Dev Biol 6: 62

Saurin AT, van der Waal MS, Medema RH, Lens SM, Kops GJ (2011) Aurora B potentiates Mps1 activation to ensure rapid checkpoint establishment at the onset of mitosis. Nature communications 2: 316

Shepperd LA, Meadows JC, Sochaj AM, Lancaster TC, Zou J, Buttrick GJ, Rappsilber J, Hardwick KG, Millar JB (2012) Phosphodependent recruitment of Bub1 and Bub3 to Spc7/KNL1 by Mph1 kinase maintains the spindle checkpoint. Curr Biol 22: 891–899

Shrestha RL, Conti D, Tamura N, Braun D, Ramalingam RA, Cieslinski K, Ries J, Draviam VM (2017) Aurora-B kinase pathway controls the lateral to end-on conversion of kinetochore-microtubule attachments in human cells. Nature communications 8: 150

Smith RJ, Cordeiro MH, Davey NE, Vallardi G, Ciliberto A, Gross F, Saurin AT (2019) PP1 and PP2A Use Opposite Phospho-dependencies to Control Distinct Processes at the Kinetochore. Cell reports 28: 2206–2219 e2208

Suijkerbuijk SJ, Vleugel M, Teixeira A, Kops GJ (2012) Integration of kinase and phosphatase activities by BUBR1 ensures formation of stable kinetochore-microtubule attachments. Developmental cell 23: 745–755

Tighe A, Staples O, Taylor S (2008) Mps1 kinase activity restrains anaphase during an unperturbed mitosis and targets Mad2 to kinetochores. J Cell Biol 181: 893–901

Tromer E, Bade D, Snel B, Kops GJ (2016) Phylogenomics-guided discovery of a novel conserved cassette of short linear motifs in BubR1 essential for the spindle checkpoint. Open Biol 6

Tromer E, Snel B, Kops GJ (2015) Widespread Recurrent Patterns of Rapid Repeat Evolution in the Kinetochore Scaffold KNL1. Genome Biol Evol 7: 2383–2393

Vallardi G, Cordeiro MH, Saurin AT (2017) A Kinase-Phosphatase Network that Regulates Kinetochore-Microtubule Attachments and the SAC. Prog Mol Subcell Biol 56: 457–484

Vleugel M, Omerzu M, Groenewold V, Hadders MA, Lens SMA, Kops G (2015) Sequential multisite phospho-regulation of KNL1-BUB3 interfaces at mitotic kinetochores. Molecular cell 57: 824–835

Vleugel M, Tromer E, Omerzu M, Groenewold V, Nijenhuis W, Snel B, Kops GJ (2013) Arrayed BUB recruitment modules in the kinetochore scaffold KNL1 promote accurate chromosome segregation. J Cell Biol 203: 943–955

Wang J, Wang Z, Yu T, Yang H, Virshup DM, Kops GJ, Lee SH, Zhou W, Li X, Xu W et al (2016a) Crystal structure of a PP2A B56-BubR1 complex and its implications for PP2A substrate recruitment and localization. Protein Cell 7: 516–526

Wang X, Bajaj R, Bollen M, Peti W, Page R (2016b) Expanding the PP2A Interactome by Defining a B56-Specific SLiM. Structure 24: 2174–2181

Waterhouse AM, Procter JB, Martin DM, Clamp M, Barton GJ (2009) Jalview Version 2--a multiple sequence alignment editor and analysis workbench. Bioinformatics (Oxford, England) 25: 1189–1191

Wijshake T, Malureanu LA, Baker DJ, Jeganathan KB, van de Sluis B, van Deursen JM (2012) Reduced life-and healthspan in mice carrying a mono-allelic BubR1 MVA mutation. PLoS Genet 8: e1003138

Wimbish RT, DeLuca JG (2020) Hec1/Ndc80 Tail Domain Function at the Kinetochore-Microtubule Interface. Front Cell Dev Biol 8: 43

Xu P, Raetz EA, Kitagawa M, Virshup DM, Lee SH (2013) BUBR1 recruits PP2A via the B56 family of targeting subunits to promote chromosome congression. Biol Open 2: 479–486

Yamagishi Y, Yang CH, Tanno Y, Watanabe Y (2012) MPS1/Mph1 phosphorylates the kinetochore protein KNL1/Spc7 to recruit SAC components. Nat Cell Biol 14: 746–752

Yang Z, Jun H, Choi CI, Yoo KH, Cho CH, Hussaini SMQ, Simmons AJ, Kim S, van Deursen JM, Baker DJ et al (2017) Age-related decline in BubR1 impairs adult hippocampal neurogenesis. Aging Cell 16: 598–601

Zhang G, Lischetti T, Nilsson J (2014) A minimal number of MELT repeats supports all the functions of KNL1 in chromosome segregation. J Cell Sci 127: 871–884

