## Supplementary Figures 1-5 for "A bifunctional kinase-phosphatase module integrates mitotic checkpoint and error-correction signalling to ensure mitotic fidelity"

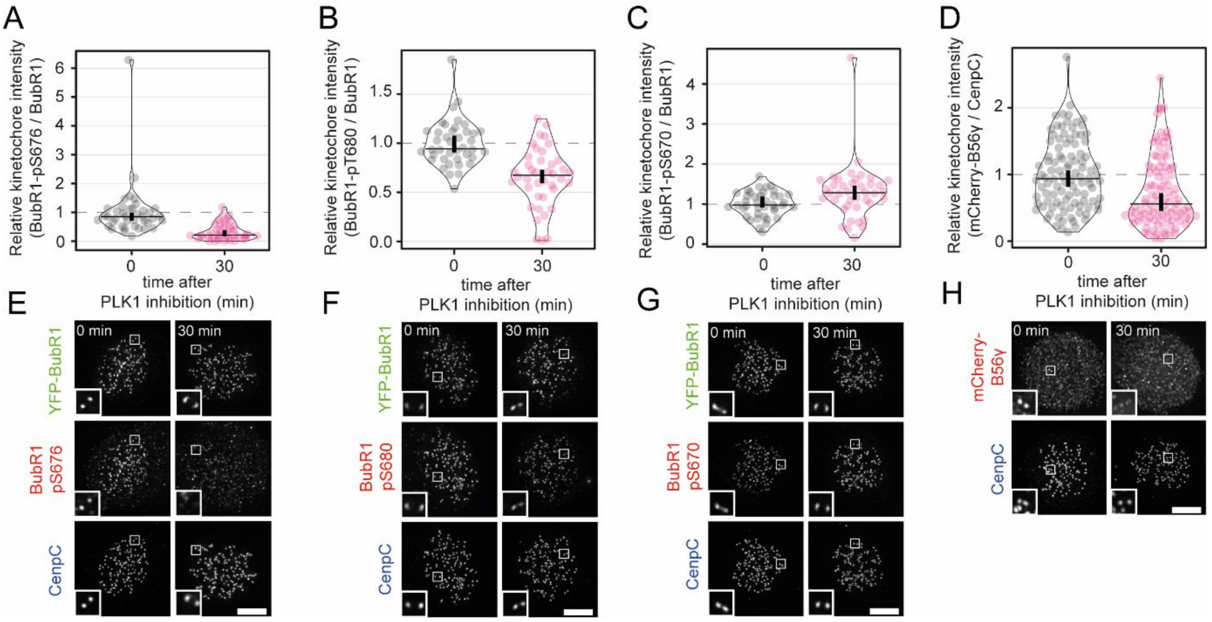


**Supplementary Figure 1 (related to Figure 1): Effect of PLK1 inhibition on PP2A-B56 recruitment sites and kinetochore localisation**

**(A-D)** Effects of PLK1 inhibition on levels of BUBR1-pS676 (A), BUBR1-pT680 (B), BUBR1-pS670 (C) and mCherry-B56γ (D) at unattached kinetochores, in nocodazole-arrested HeLa FRT cells untreated or treated with the PLK1 inhibitor BI-2536 (100 nM). Kinetochore intensities from 40-100 cells, 4 to 5 experiments. **(E-H)** Example immunofluorescence images of the kinetochore quantifications shown in 1A-D. The insets show magnifications of the outlined regions. Scale bars: 5 μm. Inset size: 1.5 μm.

Kinetochore intensities are normalised to the timepoint 0'. Violin plots show the distributions of kinetochore intensities. For each violin plot, each dot represents an individual cell, the horizontal line represents the median, while the vertical one the 95% CI of the median, which can be used for statistical comparison of different conditions (see Materials and Methods).

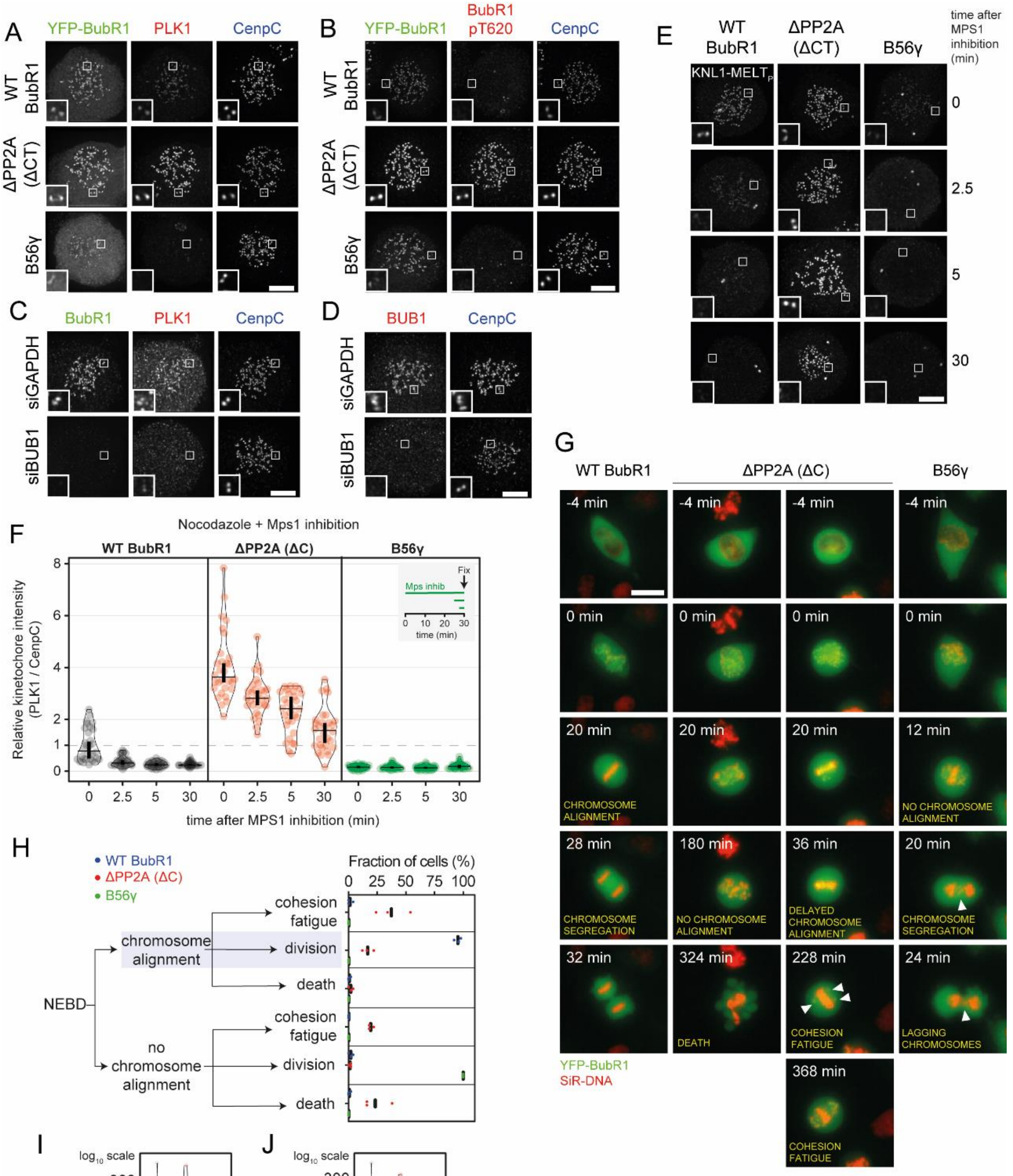


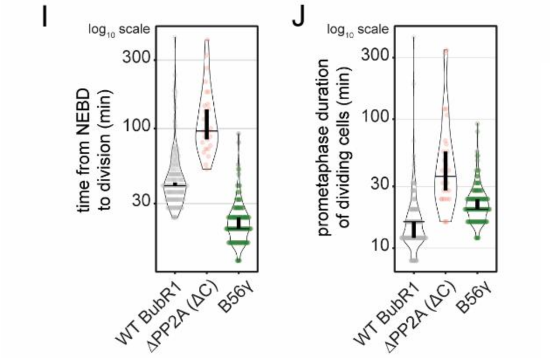


**Supplementary Figure 2 (related to Figure 2): Effect of lock PLK1 or PP2A on BUBR1, on chromosome segregation.**

**(A-E)** Example immunofluorescence images of the kinetochore quantifications shown in Figure 2B-D. (D) shows the knockdown efficiency of BUB1 – in relation to Figure 2B. The insets show magnifications of the outlined regions. Scale bars: 5 µm. Inset size: 1.5 µm. **(F)** Effects of locking PLK1 or PP2A on BUBR1 on the levels of PLK1 at unattached kinetochores, in nocodazole-arrested HeLa FRT cells expressing the indicated BUBR1 and treated with the MPS1 inhibitor AZ-3146 (2.5µM). Treatment with MG132 was included in to prevent mitotic exit after the addition of the MPS1 inhibitor. Kinetochore intensities from 30 cells, 3 experiments. **(G)** Example images from live movies highlighting the most frequent cell fates shown in Figure 2I-J. Cell fates are reported in yellow, and white arrows highlight defects during chromosome alignment or segregation. Scale bar: 20 µm. **(H)** Percentages of the cell fates shown in Figure 2I from the 3 repeats of the experiment. For each distribution, the thick line corresponds to the mean values reported in Figure 2I. **(I-J)** Duration of mitosis (I) and prometaphase (J) of cells from Figure 2I that divide after NEBD. Sample sizes: 146 cells in BubR1 WT, 27 cells in ΔPP2A (ΔC), 150 cells in B56γ. Data from 3 experiments.

Kinetochore intensities in (F) are normalised to the WT BUBR1 0' condition. Violin plots show the distributions of kinetochore intensities of PLK1 (F) or the distributions of the mitotic and prometaphase durations in a logarithmic scale (I-J). For each violin plot, each dot represents an individual cell, the horizontal line represents the median, while the vertical one the 95% CI of the median, which can be used for statistical comparison of different conditions (see Materials and Methods).

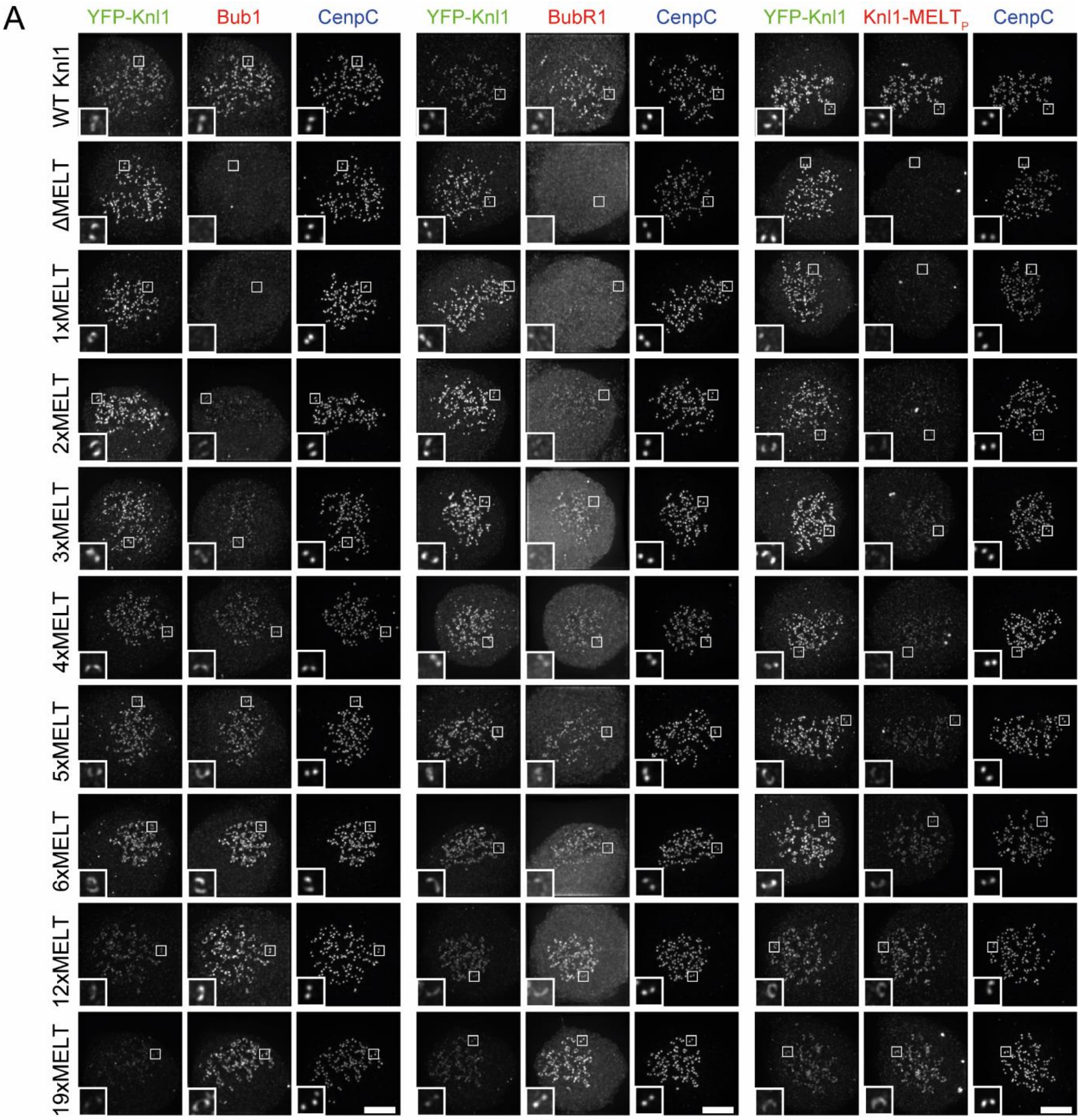


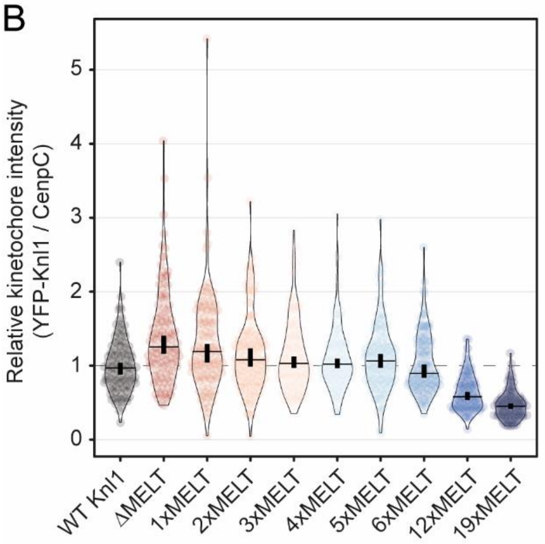


**Supplementary Figure 3 (related to Figure 3): Recruitment of KNL1, KNL1-pMELT, BUB1 and BUBR1 at kinetochores in the KNL1 MELT mutants**

**(A)** Example immunofluorescence images from the quantifications shown in Figure 3D. The insets show magnifications of the outlined regions. Scale bars: 5 µm. Inset size: 1.5 µm. **(B)** Levels of YFP-Knl1 at unattached kinetochores relative to CenpC, in nocodazole-arrested HeLa FRT cells expressing the indicated KNL1 MELT mutants described in Figure 3C. Kinetochore intensities from 120 cells, 10 experiments. Kinetochore intensities in are normalised to the WT Knl1 condition. Violin plots show the distributions of kinetochore intensities. For each violin plot, each dot represents an individual cell, the horizontal line represents the median, while the vertical one the 95% CI of the median, which can be used for statistical comparison of different conditions (see Materials and Methods).

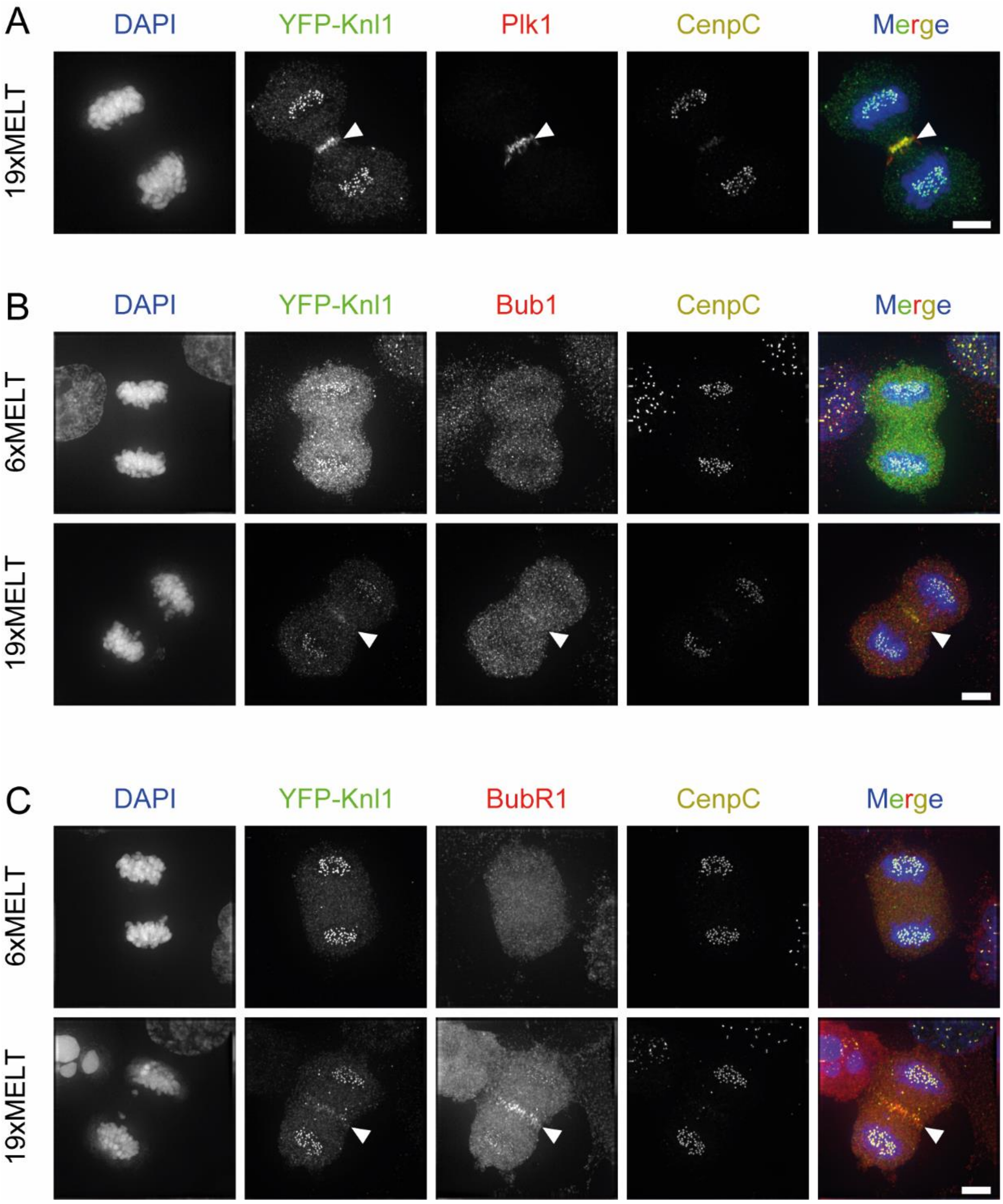


**Supplementary Figure 4 (related to Figure 4): Abnormal recruitment of KNL1, BUB1 and BUBR1 to the midbody during anaphase in KNL1-19xMELT cells**

**(A-C)** Example immunofluorescence images of 19xMELT KNL1 cells recruiting KNL1, PLK1 (A), BUB1 (B) and BUBR1 (C) at the midbody in anaphase cells. White arrows highlight the midbody. Scale bars: 5 μm.

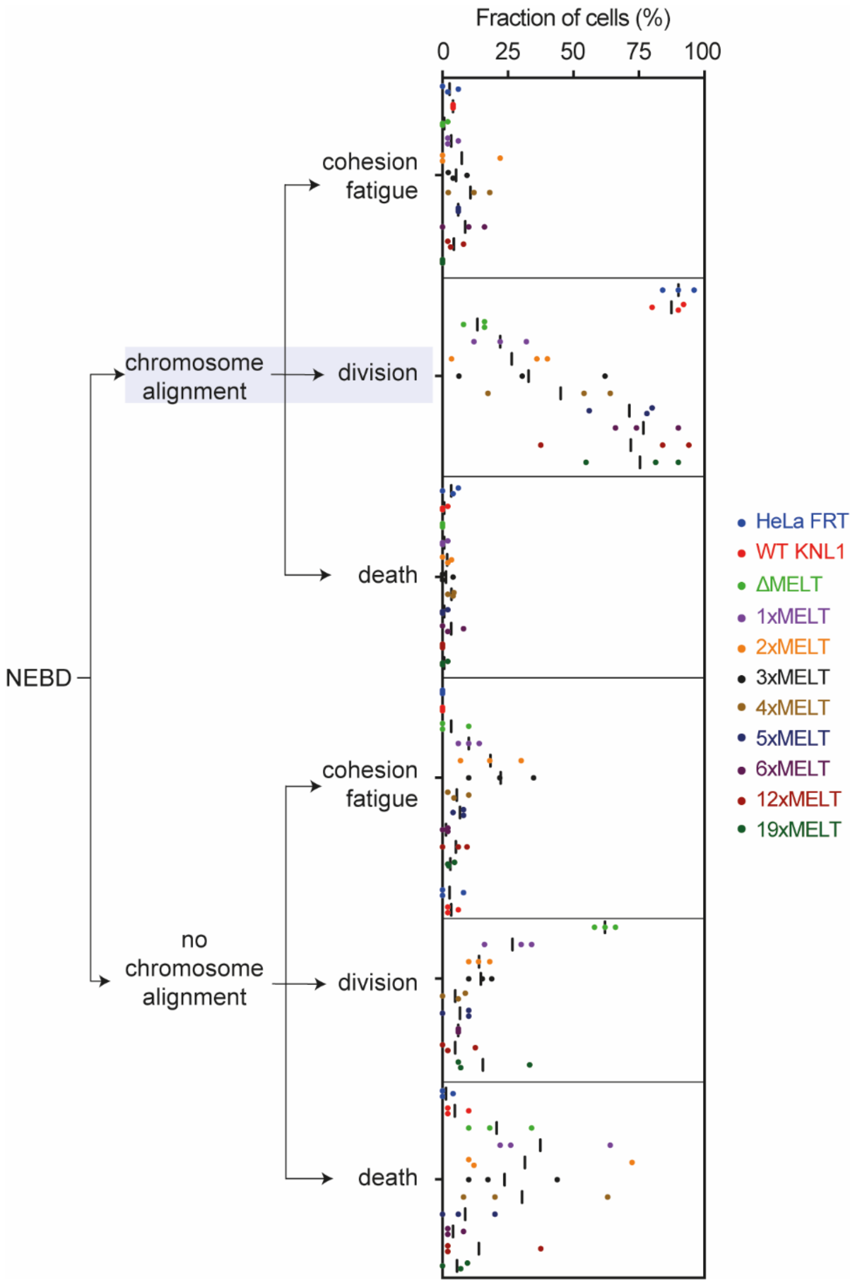


**Supplementary Figure 5 (related to Figure 5): Percentages of the different cell fates in the KNL1 MELT mutants.**

Percentages of the cell fates shown in Figure 5A from the 3 repeats of the experiment. For each distribution, the thick line corresponds to the mean values reported in Figure 5A.
